# Brain transcriptomic signatures for mood disorders and suicide phenotypes: an anterior insula and subgenual ACC network postmortem study

**DOI:** 10.1101/2024.08.14.606080

**Authors:** Dhivya Arasappan, Abigail Spears, Simran Shah, Roy D Mayfield, Nirmala Akula, Francis J. McMahon, Mbemba Jabbi

## Abstract

Mood disorders affect over ten percent of humans, but studies dissecting the brain anatomical and molecular neurobiological mechanisms underlying mood (dys)functions have not consistently identified the patterns of pathological changes in relevant brain regions. Recent studies have identified pathological changes in the anterior insula (Ant-Ins) and subgenual anterior cingulate (sgACC) brain network in mood disorders, in line with this network’s role in regulating mood/affective feeling states. Here, we applied whole-tissue RNA-sequencing measures of differentially expressed genes (DEGs) in mood disorders versus (vs.) psychiatrically unaffected controls (controls) to identify postmortem molecular pathological markers for mood disorder phenotypes. Using data-driven factor analysis of the postmortem phenotypic variables to determine relevant sources of population variances, we identified DEGs associated with mood disorder-related diagnostic phenotypes by combining gene co-expression, differential gene expression, and pathway-enrichment analyses. We found downregulation/under expression of inflammatory, and protein synthesis-related genes associated with psychiatric morbidity (i.e., all co-occurring mental disorders and suicide outcomes/death by suicide) in Ant-Ins, in contrasts to upregulation of synaptic membrane and ion channel-related genes with increased *psychiatric morbidity* in sgACC. Our results identified a preponderance of downregulated metabolic, protein synthesis, inflammatory, and synaptic membrane DEGs associated with *suicide* outcomes in relation to a factor representing *longevity* in the Ant-Ins and sgACC (AIAC) network. Our study revealed a critical brain network molecular repertoire for mood disorder phenotypes, including suicide outcomes and longevity, and provides a framework for defining dosage-sensitive (i.e., downregulated vs. upregulated) molecular signatures for mood disorder phenotypic complexity and pathological outcomes.

## INTRODUCTION

Major depressive Disorder ‘MDD’ and bipolar disorder ‘BD’, together referred to as mood disorders, are profoundly debilitating brain and behavioral disorders that globally affects about 400 million people annually. Mood disorders inflict substantial disease burden, causes nearly a million premature mortality due to suicide alone, and is associated with increased adverse socioeconomic consequences and social isolation on the global population (Kessler et al. 2005; Murray et al. 2012). The cumulative co-occurrence of mood disorders and comorbid psychiatric and chronic medical conditions like cardiovascular diseases can exert a compounding negative toll on human well-being, life expectancy, and mortality outcomes (Kessler et al. 2005; Oquendo et al. 2010; Whiteford et al. 2013; Baxter et al. 2016; Amare et al. 2017; Niculescu et al. 2017; Turecki et al. 2019). Although previous research has identified pathobiological markers for prevalent conditions like cardiovascular diseases (Gibbs et al. 2001), which often co-occur with mood disorders, the molecular neurobiological mechanisms underlying mood disorders and comorbid conditions remain unclear.

Of relevance to the behavioral ability to regulate mood functions in health and diseases, the anterior insula cortex (Ant-Ins) and subgenual anterior cingulate cortex (sgACC) brain network is well-documented to harbor the most hardwired connections with other brain regions via cortical and sub-cortical telencephalic brain connective fibers (Mesulam 1998), and with peripheral cardiovascular, gut, and adrenal systems through intricate descending extratelencephalic fibers (Levinthal & Strick 2020). The anatomical and functional integrity of the Ant-Ins aspect of this brain network is critical in engendering *<u>interoceptive</u>* sensing of internal feeling states like pain, itch, taste, smell, body temperature, mood, experience of sickness, and emotions/affect (Jabbi et al. 2008; Craig AD 2009; Deng H et al. 2022; Khalsa et al. 2018). Furthermore, the sgACC and adjacent medial prefrontal cortex aspect of this brain network translates *<u>interoceptive to exteroceptive</u>* domains by integrating external percepts like visual, auditory, olfactory, tactile, and gustatory/chemosensory cues with bodily feeling states to thereby regulate/engender emotional experiences and mood states (Nauta 1971; Goldman-Rakic 1988; Harrison et al. 2009; Joyce et al. 2020; Wang et al. 2021). Together, the anterior insula-subgenual cingulate cortical brain network (AIAC) integrates incoming stimuli with resulting feeling states induced by those stimuli or the imagination of those stimuli (Jabbi et al. 2008) and thereby codes contextually meaningful percepts (e.g., an advancing canine can potentially inflict a painful bite that can be deadly). Through this integration of perceptual and affective states, the AIAC network is hypothesized to be a critical brain system for regulating emotions and mood tone (Savitz & Drevets 2009; Rive et al. 2013).

The AIAC network is strategically situated at the cortical convergence zone of hardwired brain-body interconnected units as if this network interconnects the phylogenetically new cortical systems with the phylogenetically older subcortex and peripheral body systems (Mesulam 1998; Levinthal & Strick 2020; Joyce et al. 2020). This hardwired brain-body anatomical connective attribute makes the AIAC network a critical regulatory node for mediating mood functions (Wang et al. 2021; Strigo & Craig 2016), ranging from imagined (Jabbi et al. 2008) to actual feeling states (Meier et al. 2016). Thus, the AIAC network’s anatomical connection with peripheral systems and its critical regulatory involvement in mood states, coupled with this network’s documented anatomical changes (e.g., reduced gray matter integrity) in mood disorders (Goodkind et al. 2015; Wise et al. 2017; Jabbi et al. 2020a), underscores the potential importance of this cortical network in healthy and disease states. However, the molecular correlates of the AIAC networks involvement in coding the physiological conditions in health and disease remains unknown. We, therefore, studied the AIAC brain network transcriptional correlates for lifetime mental health, physical health and mortality outcomes in donor samples with a) no lifetime history of psychiatric illness (neurotypical controls) and in those with b) a lifetime history of mood disorders and comorbid psychiatric and medical conditions. We applied a data-reduction quantification of the AIAC network’s DEGs to test the hypothesis that this brain network’s molecular repertoire will correlate with mood disorder and complex comorbid disease phenotypes.

## METHODS

### Participants

This study was approved by the Human Brain Collection Core Oversight Committee. Clinical information on the AIAC samples are as follows: Ant-Ins samples included 100 donors, of which 37 BD, 30 MDD, and 33 unaffected controls; and sgACC samples included 152 donors, of which 38 BD, 54 MDD, and 60 unaffected controls. In addition, RNA samples were extracted from the AIAC sub-regional *postmortem* tissue (identified to be reduced in volume in brain studies of mood disorders) using a standardized procedure by the NIMH Human Brain Collection Core (HBCC).

### Brain Dissection, RNA-Extraction, and Sequencing

The NIMH Human Brain Collection Core (HBCC) provided the *postmortem* samples for which informed consent is acquired according to NIH IRB guidelines. Clinical characterization, neuropathology screening, and toxicology analyses followed previous protocols (Lipska et al. 2006). The region of interest targeted for dissection of the Ant-Ins was defined as the most anterior portion of the insula encompassing the identified reduced gram matter volume (GMV) in the completed meta-analysis by the authors (Jabbi et al. 2020a). Therefore, the dissected regional volume corresponded to the anterior portion of the Ant-Ins, where the caudate and putamen are approximately equal in size (see Supplementary Figure 1 “**Fig S1A**). Frozen tissue was dissected from the Ant-Ins section for each donor for RNA sequencing. The dissected regional volume from the sgACC was defined as a portion of the ACC Brodmann areas 32/25 (**Fig S1B**) (Akula et al. 2021).

### RNA-Extraction of Ant-Ins & sgACC

The HBCC further pulverized all dissected tissues separately and aliquoted 50mg from each sample for standardized total RNA processing. Specifically, RNeasy Lipid Tissue Mini Kit (50) was used for RNA purification using the 50 RNeasy Mini Spin Columns, Collection Tubes (1.5 ml and 2 ml), QIAzol Lysis Reagent, RNase-free Reagents, and Buffers kit from Qiagen. DNase treatment was applied to the purified RNA using a Qiagen RNase-Free DNase Set (50) kit consisting of 1500 Kunitz units RNase-free DNase I, RNase-free Buffer RDD, and RNase-free water for 50 RNA minipreps. After DNAse treatment, the purified RNA from pulverized AIAC was used separately per individual to determine RNA quality as measured in RNA integrity number (RIN) values using Agilent 6000 RNA Nano Kit consisting of the microfluidic chips, Agilent 6000 RNA Nano ladder, and reagents on Agilent 2100 Bioanalyzer. Samples with RIN < 6 were excluded from the study.

### Illumina-Sequencing, Read-Mapping, and Gene-Quantification of AIAC network

For the Ant-Ins samples, total RNA was extracted, and only samples with RNA integrity numbers (RIN values) greater than 6, as confirmed using the Agilent Bioanalyzer, were used for library preparation. Ribosomal RNA was depleted using the RiboMinus Eukaryote kit from Life Technologies (Foster City, CA, USA) for RNA-Seq and confirmed using Agilent Technologies’ Bioanalyzer (Santa Clara, CA, USA). mRNA selection was completed using the Poly(A) purist kit from Thermo Fisher, and paired-end libraries with average insert sizes of 200bp were obtained using the NEBNext Ultra II Directional RNAs Library Prep kit from New England BioLabs. All 100 samples were processed and sequenced on the Illumina HiSeq 4000 at the Genome Sequencing and Analysis Facility (GSAF: https://wikis.utexas.edu/display/GSAF/Home+Page) at UT Austin, USA (Supplementary Methods). Thirty million paired end reads per sample (150 base pairs in length) were generated by sequencing runs of 4 samples per lane of the sequencer. First, sequenced reads were assessed for quality with Fastqc to assess sequencing reads for median base quality, average base quality, sequence duplication, over-represented sequences, and adapter contamination (Andrews 2010). We looked at median base quality, average base quality, sequence duplication, over-represented sequences, and adapter contamination, which were < 5%. Median quality at every base was > 30 for all samples, and more than 90% of the reads had average base quality > 30, making read trimming or filtering redundant. We did not remove samples because all samples had typical sequence duplications < 60%, as in high-coverage RNA-Seq data, and no adaptor trimming was performed (i.e., adaptor contamination percentages < 5%). Next, the reads were pseudo-aligned to the human reference transcriptome (GRCh38-encode) using Kallisto (Bray et al. 2016), and gene-level abundances were obtained.

For the sgACC, the RNA sequencing method and protocol were described earlier in the original study (see Akula et al. 2021). Briefly, total RNA extracted from frozen dissections of sgACC and only samples with RNA integrity numbers (RIN values) greater than 6, as confirmed using the Agilent Bioanalyzer, were used for library preparation. Total RNA was captured using the RiboZero protocol, followed by library preparation, and stranded paired-end sequencing was performed on the RNA samples using the Illumina HiSeq 2500 system. We obtained an average of two hundred and seventy million reads per sample, totaling ∼54 billion reads. After quality control, reads were mapped to human genome build 38 using Hisat2 (Pertea et al. 2016). Finally, gene and transcript counts were obtained using StringTie (Pertea et al. 2016).

For both Ant-Ins and sgACC, any genes expressing 0 in 80% or more samples were filtered out to remove low-count genes from further analysis. Next, the abundances were normalized using DESeq2 and transformed with variance stabilizing transformation (a transformation to yield counts that are approximately homoscedastic, having a constant variance regardless of the mean expression value). Finally, Principal Component Analysis was performed using 25% of the highest variance genes to explore the underlying data’s structure and the largest sources of variance. Lastly, genes with an expression value of 0 in 80% of samples or more were removed from further analysis to correct for sporadically large fold-change outliers.

### Statistical Analysis

#### Weighted Gene Co-Expression Network Analysis (WGCNA)

Scale-free co-expression networks were constructed with gene abundances using the WGCNA package in R (Langfelder & Horvath 2008) (See **Fig 1** for data analytics workflow). WGCNA provides a global perspective and allows the identification of co-expressed gene modules. It avoids relying on arbitrary cut-offs involved in selecting differentially expressed genes. Instead, it identifies a group of genes changing in the same direction and magnitude, even if these changes are smaller. WGCNA identifies co-expressed modules of genes, thereby identifying genes that are likely co-regulated or may belong to the same functional pathway, using a dynamic tree-cutting algorithm based on hierarchical clustering (i.e., minimum module size=30). A given module’s eigengene, defined as the first principal component of the expression matrix of the corresponding module, can be correlated to sample variables to identify modules of interest. We correlated the module eigengenes to different postmortem sample characteristics and selected the two modules that showed significant correlation to variables of interest, such as *diagnostic and suicide-linked variables*. Driver genes (i.e., genes within co-expressed gene modules whose distinct expression patterns are similar to the overall expression profile of the entire co-expressed modules) were used to identify pathobiological functions associated with each module.

**Figure 1.**
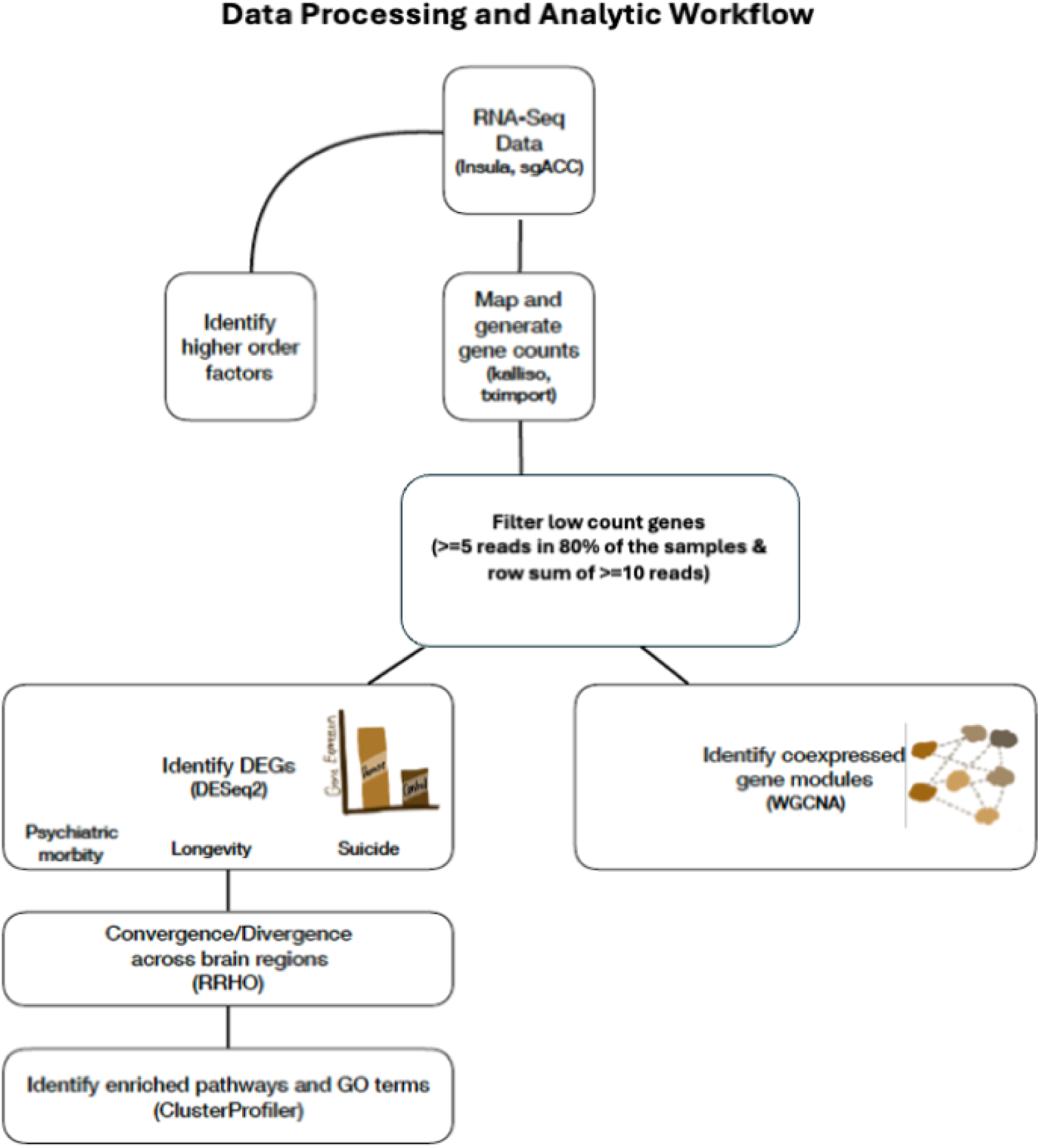
Analytic Workflow. From top to bottom, the workflow illustrates the step-by-step processing of RNA-seq data and analysis in the AIAC network donors including the 180 unique samples.

#### *Postmortem* variable factor-analysis

The *postmortem* variables included *mood disorder diagnoses; # of lifetime-Axis-I diagnostic occurrences* (e.g., Axis-I-loading of the number of comorbid disorders such as (poly)-substance use disorders, psychosis, anxiety, eating disorders, etc.); # of *lifetime-Axis-III diagnoses* (e.g., medical conditions such as diabetes, cancer, cardiovascular disease, etc.)*; manner of death* (e.g., natural, suicides, homicides or accidents) and *cause of death* as specified by the medical examiner reports (e.g., blunt force trauma to the chest, gunshot, motor vehicle accident, drowning, hanging, etc.)*; demographics* (race, age at death, sex, years of education, number of children/fecundity, and marital records); *technical variables* (brain-weight, postmortem interval, pH, and RIN-values); and *toxicology* (blood alcohol/blood narcotics levels). We applied Principal Axis Factoring using the Oblimin Rotation with Kaizer Normalization (Costello & Osborne 2005) to identify higher-order factors explaining the differences in *postmortem* variables. We also included those with commonalities of ≥ 0.45.

#### Differential Gene Expression Analysis

We first compared gene expression profiles with the two regional datasets by conducting simple comparisons across MDD vs. controls and bipolar disorder vs. controls separately.

Our exploratory factor analysis assessed the relationship between all the postmortem variables to determine the existence of higher-order factor loadings that better explain postmortem variance than the original variables (see results section for full details). In order to remove low signal or outlier genes from the differential expression analysis, two filters were used. <u>Only genes with 5 reads or more in 80% of the samples and a row sum of 10 reads or more</u> <u>were considered for further analysis</u>. These filters help prevent genes which are undetected in most samples and lowly expressed in a few samples from being identified as differentially expressed genes. To identify gene expression signatures related to differences in our identified higher-order factors, such as *psychiatric morbidity*, we compared high (samples scoring above the median split of the factor loading) scores on *psychiatric morbidity* vs. low (samples scoring below the median split of the factor loadings) scores of this factor. Similar comparisons were carried out for *longevity*. Differential gene expression between samples differing in *psychiatric morbidity* and *longevity* status was assessed across the AIAC network based on the negative binomial distribution for modeled gene counts using DESeq2 (Anders & Huber 2010). In addition, RIN-values were included in the DESeq2 design matrix as a covariate to control for potential confounds.

Controls were omitted in the last comparison (i.e., to examine gene expression profiles that might be linked explicitly to suicide completion vs. non-suicide deaths in persons diagnosed with mood and comorbid psychiatric disorders. Therefore, only genes with corrected p-value (after Benjamini-Hochberg multiple testing corrections) ≤ 0.05 are reported as significantly differentially expressed. GO-terms enriched in these genes were identified using Enrichr (Chen et al., 2013; Kuleshov et al., 2016).

#### AIAC network Rank Rank Hypergeometric Overlap (RRHO) analysis

We applied the stratified RRHO method implemented by Cahill et al. (Cahill et al. 2018), an updated and advanced version of previous applications of RRHO using R (Cahill et al. 2018). In essence, the updated RRHO algorithm is designed to quantify the preponderance (significance) of correlation or overlap between two gene lists from two sets of independent experiments or datasets like our examples of the datasets from the AIAC network in the current study in terms of upregulation or downregulation based on enrichment measures. The updated RRHO algorithm, or “Stratified method,” calculates the degree of overlap based on quadrant-specific analyses (see Figure 6A-C) (Cahill et al. 2018). Precisely, the updated method designed a new approach that takes each quadrant and counts from the outward corner to the cutoff point to define the number of genes from the first gene expression dataset (Ant-Ins) and the number of genes from the second dataset (sgACC) and assess the overlapping enrichment between them.

## RESULTS

### Demographics, morbidity and mortality variability, and global DEGs across samples

Overall, 100 donors with dissected brain tissue and successful RNA sample extraction from the Ant-Ins region were included in the study: 33 psychiatrically unaffected controls/controls (0 suicide), 37 BD (28 suicide), and 30 MDD (24 suicide) donors. For the sgACC region, 152 samples were included in the study: 60 controls (0 suicide), 38 BD (28 suicide), and 54 MDD (42 suicide) donors. Notably, of the 180 unique donors, 72 of them were brain donors with both Ant-Ins and sgACC extracted RNA samples included in the current study.

To first examine the degree of mood disorder co-occurrence with other psychiatric (Axis-I) and medical (Axis-III) conditions, we used an analysis of variance (ANOVA) to assess if the presence of chronic medical conditions like cardiovascular diseases, cancers, and diabetes differ between BD, MDD, and controls. We found that comorbidity with chronic medical conditions was highest in mood disorders (at F=5.72, p=0.004) and more so in MDD vs. controls, followed by bipolar disorder vs. controls, even though a proportion of controls died from terminal Axis-III conditions (**Table 1**). We then assessed the degree of Axis I comorbidities like having lifetime BD with co-occuring anxiety, polysubstance use, psychosis in the same donor; or having lifetime MDD with co-occuring psychosis, anxiety, post-traumatic stress disorder, alcohol use disorder all in same donor (degree of psychiatric comorbidity across the samples) and found no differences between the MDD and BD samples looking at both the Ant-Ins and sgACC region donors. We further evaluated postmortem body mass index (BMI) differences across all samples and found no association between diagnoses (mood disorders vs. controls) and BMI in the overall Ant-Ins samples. However, the total sgACC samples (including 72% of the Ant-Ins samples) showed increased BMI in unaffected controls compared with the mood disorder donors (F=3.7, p=0.027).

**TABLE 1.** Psychiatric and Chronic Disease/Medical Comorbidity.

| **TABLE 1 Psychiatric and Chronic Disease/Medical Comorbidity** |  |  |  |  |  |  |  |  |  |
| --- | --- | --- | --- | --- | --- | --- | --- | --- | --- |
|  | % of Comorbid Axis III Diseases |  |  |  |  |  |  |  |  |
| **Primary Axis I Diagnosis** | Lung | Cardiovascular | Cancer | Diabetes/Endocr. | Inflammatory/Chronic Pain | Subst. Intoxication/Poisoning | Other CNS | Infection | # of Donors |
| Bipolar Disorder (BD) | 17.3% | 36.5% | 9.6% | 19.3% | 15.4% | 42.3% | 19.2% | 5.8% | 52 |
| Major Depressive Disorder (MDD) | 13.3% | 50.0% | 5.0% | 16.7% | 5.0% | 46.7% | 8.3% | 5.0% | 60 |
| Unaffected controls | 8.8% | 74.4% | 7.4% | 13.2% | 4.4% | 0.0% | 2.9% | 2.9% | 68 |
| Abbreviations: Lung = lung disease; Endocr. = Endocrine diseases/obesity; Subst. = Substance; CNS = Central Nervous System Diseases such as migraine or epilepsy with no focal localization in the AIAC network; # = number |  |  |  |  |  | Total # of Unique Donors |  |  | 180 |

### Weighted gene co-expression network analysis (WGCNA) identifies disease DEG modules

To assess the global gene co-expression profiles for mood disorder diagnoses, other demographics variability, psychiatric disorder and chronic medical disease comorbidity, and suicide mortality-related outcomes across the AIAC network, we performed WGCNA (Langfelder & Horvath 2008) of the two regions separately. Our WGCNA co-expression matrix underwent “soft thresholding” by restricting the number of modules to 14. Closely related co-expression modules were merged into one module based on a dissimilarity (1 – correlation) cut height of 0.4 for any of the two brain regions with more than 14 co-expression modules at 0=thresholding. We defined correlation matrixes for the following variables of interest: mood disorder diagnoses, age at death, Axis-I (comorbid psychiatric conditions, including diagnoses of each of the studied mood disorders as primary psychiatric diagnoses), Axis-III/chronic medical conditions, BMI, and measures of suicide lethality for our WGCNA model. The functionality of the related co-expression pathways was defined using the Gene Ontology (GO) toolbox to identify enriched GO terms (Kuleshov et al., 2016) for each specified WGCNA module.

We further examine gene co-expression beyond the measures of psychiatric phenotypes by assessing Axis II/chronic disease comorbidity related gene expression modules in the AIAC network (see **Fig 1** for analytic steps). We found that Axis-III comorbidity correlated negatively with the Ant-Ins <u>tan module</u>, which is enriched for metabolic, energy transport, mitochondrial translation/gene co-expression genes (**Fig 2A & supplementary figure 1A “Fig S2A”**). Age at death, Axis-I, and suicide lethality collectively correlated negatively with the <u>yellow module</u> capturing cellular and neuronal ion channel/calcium ion-dependent signaling and synaptic membrane gene co-expression (Schloss et al. 2004; Ryding et al. 2006) (**Fig 2A & Fig S2B**), and the <u>black module</u> enriched for O-glycan synthesis and inflammatory cytokine signaling gene co-expression in Ant-Ins (Miller & Raison 2016). Axis-I psychiatric comorbidity was also correlated positively with the <u>brown module</u> enriched for a wide-ranging inflammatory cytokine response, T-cell immune response and leukocyte functions gene co-expression in the Ant-Ins (**Fig 2A & Fig S3A**) and Axis-III comorbidity correlated positively with the <u>magenta/green</u> <u>module</u> enriched for biosynthesis/protein synthesis and viral transcription in Ant-Ins (**Fig 2A & Fig S3B**). Together, these WGCNA findings of enriched ion channel and inflammatory signaling in the Ant-Ins are convergently associated with comorbid lifetime physical (Axis II) and mental (Axis I/psychiatric disease), suggesting a likely general disease state-related molecular repertoire in the Ant-Ins that is beyond mood and psychiatric disorders.

**Figure 2.**
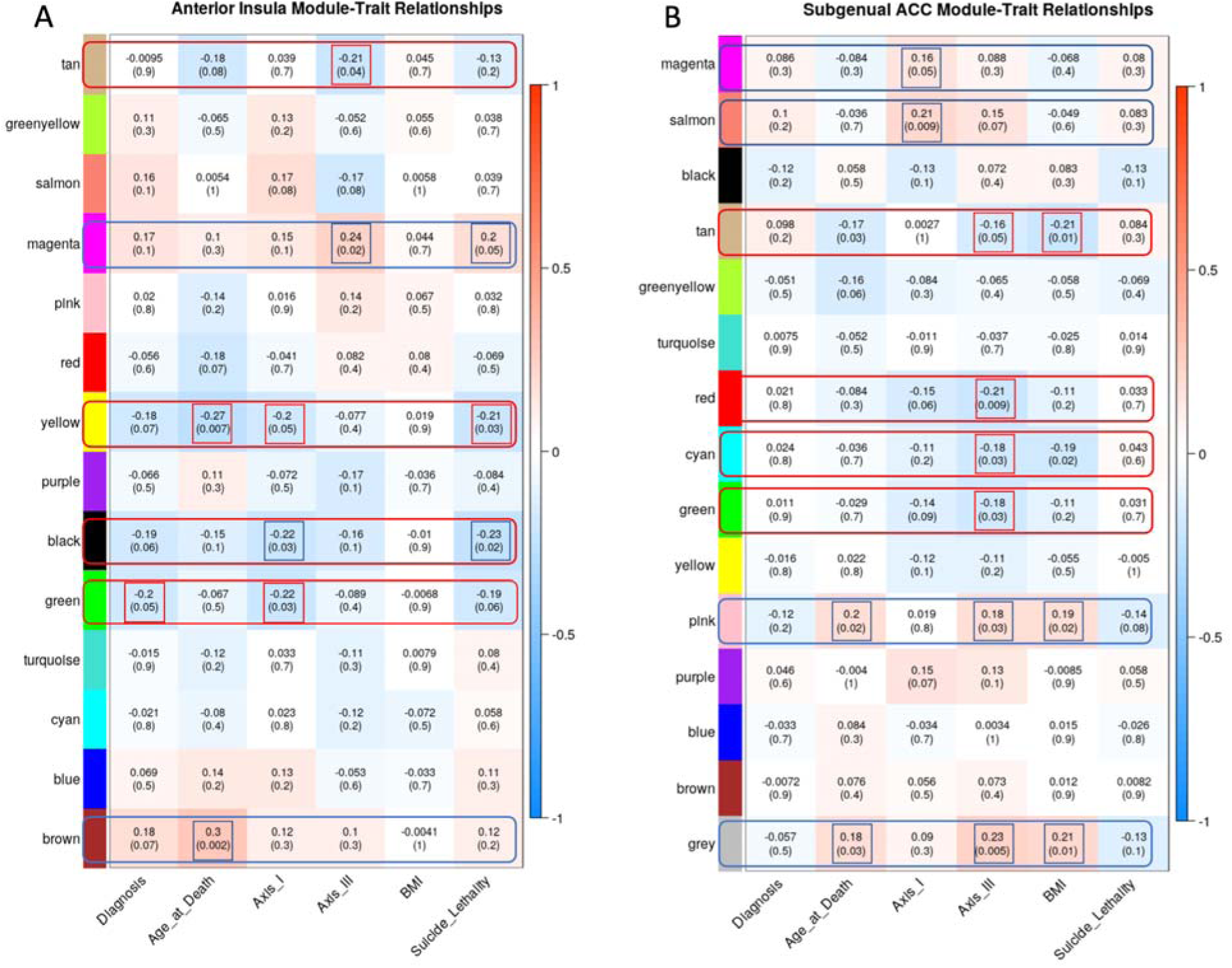
Weighted Gene Co-expression Network Analysis. **A-B**, the y-axis illustrates the WGCNA heatmaps of correlations between specific gene expression modules. The x-axis represents specific clinical phenotypes such as lifetime mental health diagnosis (Diagnosis), age at which the donors died (Age at Death), comorbid lifetime mental disorders (Axis I), comorbid lifetime physical diseases (Axis III), body mass index (BMI), and the lethality of the suicide method for those who completed suicide.

We assessed WGCNA for the sgACC data and identified a positive correlation between Axis-I and the <u>salmon module</u> enriched for spliceosome, thyroid hormone, and notch signaling gene co-expression (**Fig 2B & Fig S4A)**. On the other hand, Axis-III comorbidity and BMI correlated negatively with the <u>tan module</u> enriched for ribosomal, spliceosomal, mRNA transport and methylation, and protein synthesis (Pishva et al., 2017; Akula et al. 2021) gene co-expression in sgACC (**Fig 2B & Fig S4B)**. Furthermore, Axis-III comorbidity correlated negatively with the <u>grey module</u> capturing cellular immune and developmental regulatory gene co-expression in the sgACC (**Fig S5A**). The <u>red,</u> pink, <u>cyan, tan, grey and green modules</u> known to be enriched for metabolic, protein synthesis and bodily homeostatic regulatory gene co-expression, were also identified in sgACC in association with Axis-III comorbidity and BMI (**Fig 2B & Fig S5B**).

In summary, WGCNA of the AIAC network revealed specific and overlapping global gene co-expression repertoires associated with psychiatric comorbidity and suicide phenotypes and Axis-III and BMI-related indications. Furthermore, whereas age at death and suicide phenotypic variability showed correlations with AIAC network gene co-expression, Axis-III was found to be correlated with twice as many co-expression modules than Axis-I in the sgACC, suggesting the relevance of the AIAC brain network’s molecular integrity for peripheral disease burden. Based on these findings, we conducted data reduction identification of relevant variables for further gene expression analysis (see details in the methods section).

### Mood disorder-specific differentially gene expression analysis identified DEGs

Using a statistical threshold of q=0.05 adjusted for multiple comparisons using false discovery rate correction (FDR) (Benjamini & Hochberg 1995), we assessed differential gene expression in MDD versus (vs.) controls and in BD vs. controls to identify diagnosis-specific DEGs across the Ant-Ins and the sgACC regions. Our identified DEGs in the Ant-Ins in MDD vs. controls included <u>five</u> genes, namely: a downregulated *SELE* gene known to control leukocyte regulation of inflammation, and four upregulated genes including the cytokine interleukin-1 receptor-like *IL1RL1* gene, a gene that regulates *IL-33/ST2* (Uversky 2014), a phosphorylated protein binding *FBXO47* gene, a mitochondrial electron transporter *MTCO2P12*, and a long-noncoding RNA (lncRNA) H19 in the Ant-Ins (**Table 2A**). To compare BD>controls, we found <u>one</u> lncRNA RP1-193H18.3 to be upregulated (**Table 2B**). Furthermore, we found no DEGs for MDD>controls or BD>controls in the sgACC at the adjusted p-value of 0.05 FDR.

**TABLE 2.**
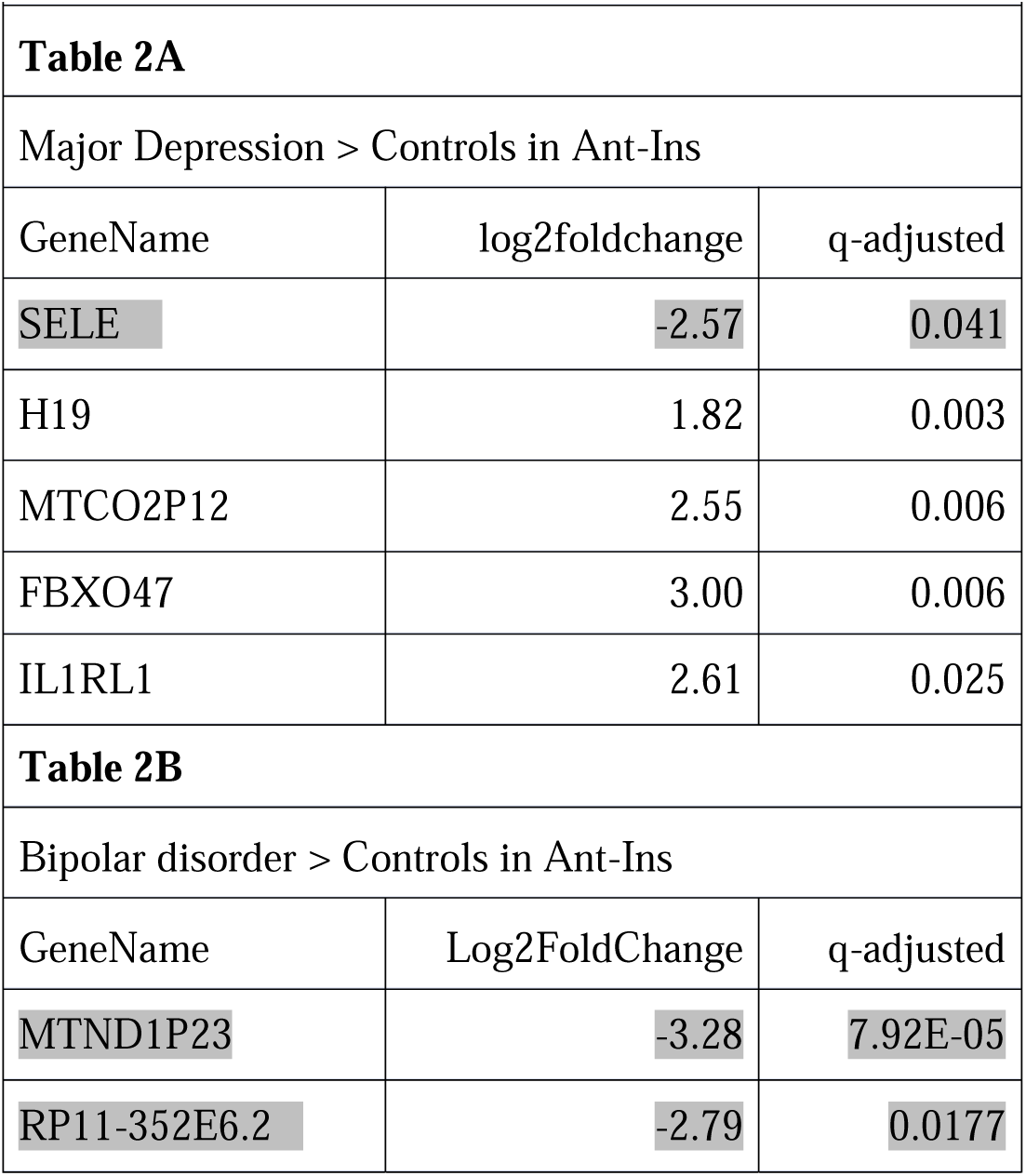
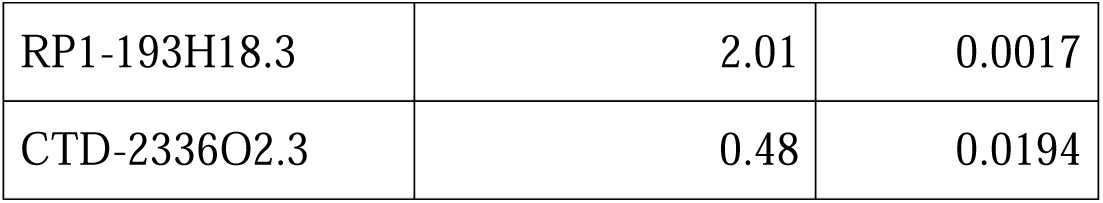

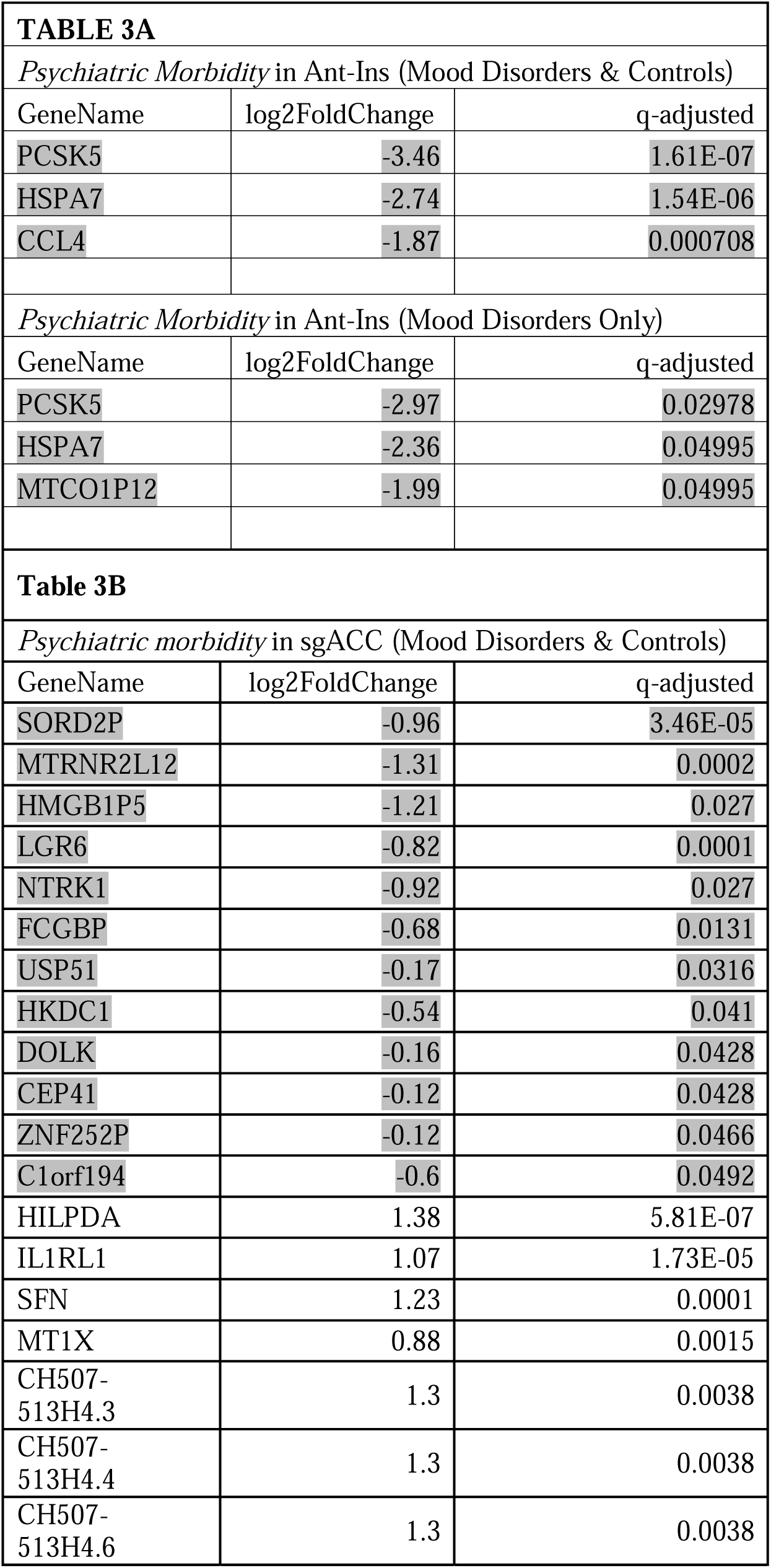
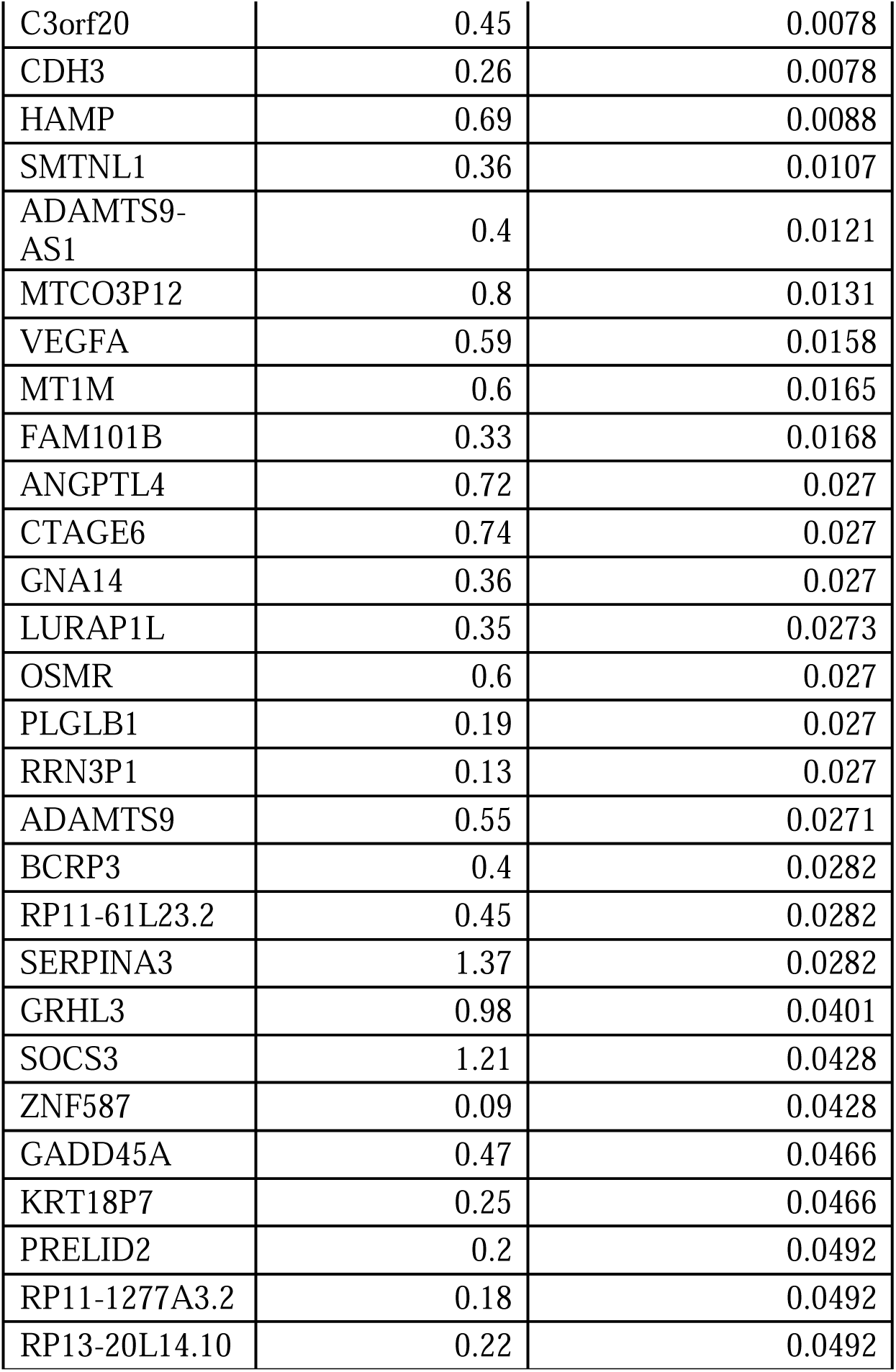

### Factor analysis identified relevant morbidity and mortality indicators

To better examine the inter-relationship between complex disease comorbidity and underlying brain molecular pathology as measured in AIAC network postmortem gene expression measures using whole tissue RNA-seq in donors who died of both chronic medical conditions (Axis-III), mood disorders related suicide, we applied a factor analytic data reduction to identify hidden phenotypic variability in our data that may influence DEGs. The application of a factor analysis of the postmortem phenotypic data is important because it allows a data-driven method of assessing what aggregate/composite variabilities could be driving biological gene expression changes (DEGs) in the studied sample without relying on predefined variables like diagnosis, age or sex which may not be sufficiently driving biological variability related to mood disorder metrics. To this aim, we included diagnoses, Axis-I, Axis-III, BMI, age at death, and suicide lethality variables, etc., in a factor analytical model using principal axis factoring for identifying higher-order variables that are more sensitive for precise quantification of phenotype-related DEGs (Jabbi et al. 2020a; Arasappan et al. 2021).

We found three higher-order factors that cumulatively explained 42.22% of the total variance of the postmortem phenotypes. These higher-order factors included *longevity/*aging (hereafter referred to as *longevity*), with 17.052% of the variance explained by this factor being found to relate to 1) <u>marital status</u> (with 0.834 factor loading), 2) <u># of children</u> (with 0.643 factor loading), 3) <u>Axis-III</u> (all non-psychiatric medical conditions) comorbidity (with 0.635 factor loading), and 4) <u>age at death</u> (with 0.584 factor loading)]. In addition, a second factor related to *psychiatric disorder morbidity and suicide outcomes* was computed, with 16.27% of the variance explained by this factor relating to 1) <u>mood disorder diagnosis</u> (with 0.897 factor loading), 2) <u>Axis-I comorbidity</u> (0.695), and 3) <u>suicide lethality</u> (0.670)]. Finally, another higher-order factor related to *RNA integrity number* (RIN) was computed, with 8.89% of the variance explained by this factor relating to 1) *<u>RIN-value</u>*(with 0.801 factor loading) and 2) <u>sex</u> (with −0.494 factor loading)]. We further quantified DEGs using our two top identified factors as higher-order factors/variables of interest. Given the importance of mood disorder as the primary psychiatric diagnostic indices of the study samples, we first analyzed the DEGs associated with *psychiatric comorbidity*, and whether the donors died by suicide to assess the transcriptomic correlates for suicide mortality, and then assessed *longevity* associated DEGs. Given that *RIN-value* and sex co-aggregated strongly, and we included RIN as a covariate to correct for potential tissue qualitative confounds and further minimize the effects of potential sex-related confounds, the RIN-Value higher-order factor was not further analyzed.

### Psychiatric (co)morbidity-related differential gene expression analysis identified DEGs

We first assessed DEGs associated with *psychiatric morbidity* (which is a measure of all co-occurring mental disorders/neuropsychiatric load and suicide outcomes). All humans, including healthy people, often undergo periodic experiences of positive and negative mood changes throughout their lifespans. Given that human mood states fluctuate between positive and negative and sometimes at extremes of these mood states, it was expected that our controls were not precluded from having lifetime sub-diagnostic threshold episodic negative mood symptoms/states. Therefore, we included the controls in our factor analysis and initial differential gene expression analyses that assessed transcript abundance associated with *psychiatric morbidity* and related phenotypes independent of diagnoses. Controls were further removed from secondary analysis to assess DEGs associated with *psychiatric morbidity* within the mood disorder samples.

Using this approach, we then applied a median split-half method of identifying DEGs associated with high vs. low psychiatric morbidity (including MDD, BD, and unaffected control samples in our analytic model) across the AIAC network at p<=0.05 FDR. In the Ant -Ins, we found <u>three</u> downregulated DEGs that recapitulated our mood disorder vs. control findings (see **Table 3**), including the protein synthesis *PSK5* gene and ATP-binding heat shock protein *HSPA7* gene (Pantazatos et al. 2017), and a mitogen-inducible monokine called C-C motif chemokine ligand-4 immunoregulatory and inflammatory *CCL4* gene (**Table 3A**; **Fig 3A-B**). We then performed a secondary analysis comparing high vs. low *psychiatric morbidity* in the mood disorder samples (excluding controls) to assess if our identified Ant-Ins DEGs are proximate to mood pathology. For this mood disorder-specific analysis, we found two of the three downregulated genes in the mood disorders and control analysis including *PSK5* and *HSPA7* as well as the mitochondrial electron transporter *MTCO2P12*, surviving p<=0.01 FDR (**Table 3A**; **Fig 3A-B**). Further, our observed high *psychiatric morbidity* GO-terms were enriched for immune/inflammatory, protein synthesis, complement activation, and Fc-gamma receptor signaling DEGs in Ant-Ins (Laguesse & Ron 2020) (**Fig 3A-B**).

**Figure 3.**
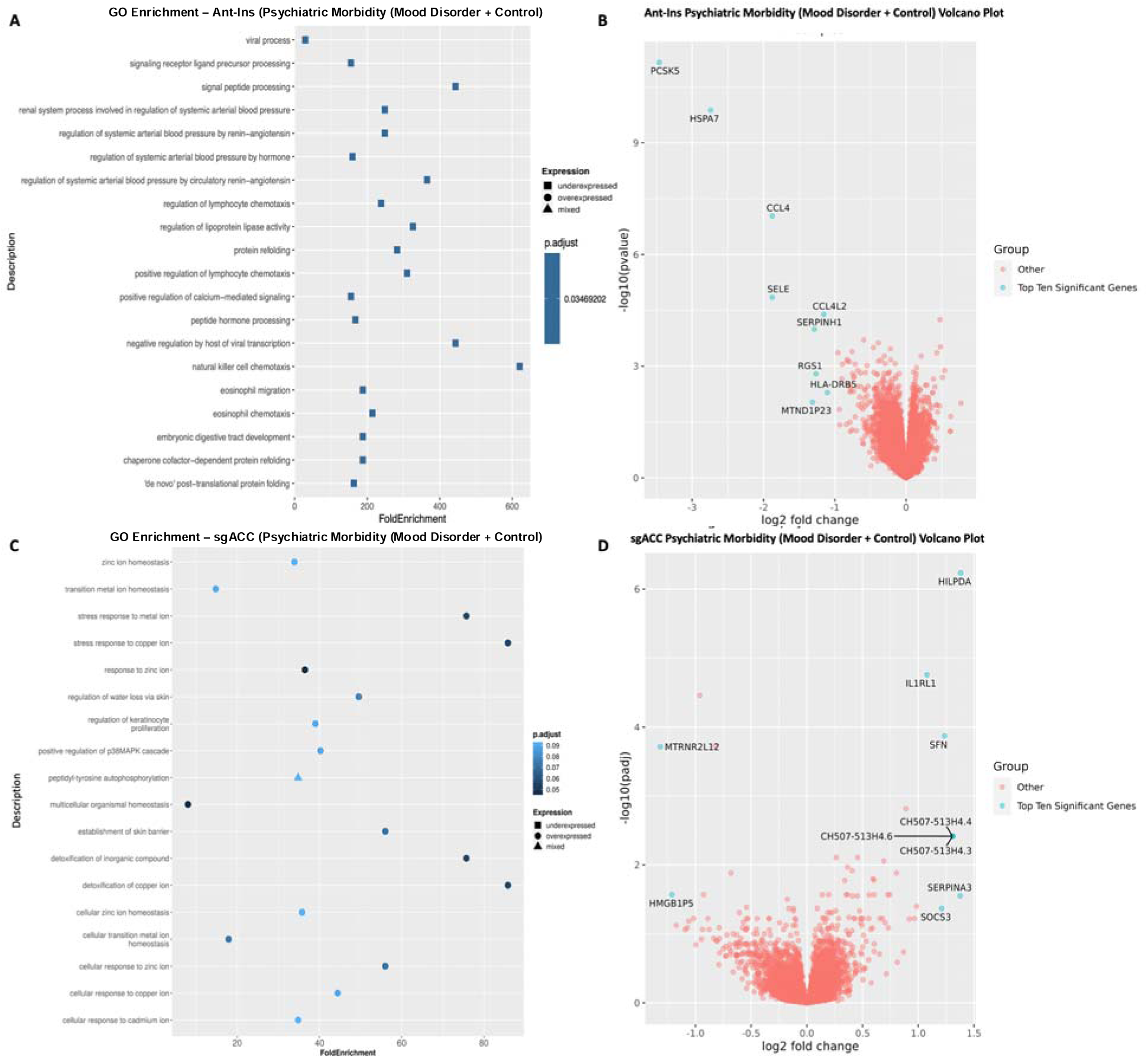
Gene Ontology (GO) terms and Volcano plots for High vs. Low Psychiatric Morbidity (higher-order factor loadings were used to compare high comorbid/morbidity versus low morbidity individuals in a differential gene expression analysis). **A**) illustrates GO terms for GECs in Ant-Ins for high vs. low psychiatric morbidity, with **B**) depicting the related volcano plot for the Ant-Ins results in **A**. **C & D**) illustrates GO terms in sgACC representing high vs. low psychiatric morbidity, and related volcano plot for the sgACC results in **C**. Because the -log10 (adjusted q-values) of a large number of genes in Insula was close to zero, volcano plots were generated using q-value instead of adjusted q-value. Genes meeting the following cutoffs: adjusted q-value <0.05 and absolute log2 fold change >= 1 were highlighted on the volcano plot as significant genes.

Given that our median split-half method included all samples, we conducted an additional Ant-Ins analysis of DEGs by comparing the subgroup of donors at the two extremes (i.e., 20 donors with the lowest scores on *psychiatric morbidity* vs. 20 donors with the highest scores on *psychiatric morbidity*). First, we compared *psychiatric morbidity* for the 20 samples with the lowest psychiatric morbidity vs. 20 with the highest psychiatric morbidity using adjusted p<0.05 FDR cut-off FDR. As a result, we found <u>one</u> downregulated DEG, namely the Neuronal PAS Domain Protein 4 master transcriptional regulator (*NPAS4)* involved in an array of biological functions, including physiological and developmental events gene (**Table S4A**). We then repeated a similar analysis comparing the 20 lowest psychiatric morbidity samples in the mood disorder cohort (excluding controls) vs. the 20 highest psychiatric morbidities in the mood disorder samples. We found no DEGs surviving p<=0.05 FDR, suggesting that Ant-Ins DEGs may not be sensitive to extreme differences in measures of *psychiatric morbidity within disease samples* (**Table S4B**).

Differential gene expression analysis of sgACC samples for high vs. low *psychiatric morbidity* in all samples including mood disorders and controls yielded forty-seven DEGs, including <u>twelve</u> downregulated differentially expressed genes (**Table 3B**; **Fig 3C-D**). DEGs for high vs. low *psychiatric morbidity* in sgACC also revealed <u>thirty five</u> upregulated genes, including the autocrine signaling lipid storage and metabolism gene *HILPDA* implicated in stress responsiveness/physical activity/energy expenditure (Cantarelli et al. 2014; Vandekopple et al. 2017; Vandekopple et al. 2019), interleukin 1 receptor-like *IL1RL1* (Uversky 2014; Kathryn et al. 2018), mitotic translational regulator *SFN*, synaptic membrane/calcium ion channel *MT1X* (Sokolowski et al. 2018; Zandi et al. 2022), an uncharacterized protein *C3orf20*, Calcium-dependent adhesion protein *CDH3* genes, the iron homeostatic hepcidin antimicrobial peptide *HAMP* gene, and the CH507-513H4.3, CH507-513H4.4, and CH507-513H4.6 novel transcripts, etc. (**Table 3B**; **Fig 3C-D**). Of interest, replicating the high vs. low *psychiatric morbidity* comparison in the mood disorder samples only (excluding controls) yielded no DEGs in the sgACC, as if the sgACC gene regulatory repertoire likely underpins the presence of mood disorder diagnosis rather than the graded disease morbidity or severity. On the other hand, the GO-terms for *psychiatric morbidity-associated* DEGs in all samples of the sgACC identified enriched pathways for cellular signaling, zinc ion homeostasis, multicellular organismal homeostasis, and metabolic balance (Zhang et al. 2022) (**Fig 3C-D**). Notably, although the unaffected controls have no recorded mental disorder history, including them in the high vs. low *psychiatric morbidity* analysis for AIAC network did not dampen the number of DEGs in the studied network.

Similar to our Ant-Ins analysis of comparing extreme scores for *psychiatric morbidity*, we assessed the sgACC transcriptome measures of DEGs by comparing 20 donors with the lowest scores on *psychiatric morbidity* vs. 20 donors with the highest scores on *psychiatric morbidity* by first including both the mood disorder and control cohorts at p<0.05 FDR. As a result, we <u>found</u> <u>five</u> downregulated and <u>thirty-three</u> upregulated DEGs (**Table S5A**). Then, we repeated this analysis by excluding the controls and only comparing the 20 lowest *psychiatric morbidity* scores within the mood disorder cohort vs. the 20 highest *psychiatric morbidity* scores and found <u>eighteen</u> downregulated and 400 upregulated DEGs (**Table S5B**).

Our additional comparison of *psychiatric morbidity* scores in the lowest and highest extremes, in the totality of all samples, and comparing the 20 lowest vs. the 20 highest psychiatric morbidity scores in mood disorder samples alone resulted in more DEGs in the sgACC, unlike the Ant-Ins that did not show any highest 20 vs. lowest 20 scoring *psychiatric morbidity* related DEGs. Together, these findings suggest that extreme comparisons may reveal related and unique molecular profiles that are differentially mediated in different brain network nodes.

At the global level, we found more differentially expressed genes (DEGs) in sgACC region compared to Ant-Ins, when using psychiatric comorbidity when comparing gene expression in suicide completer vs. non suicide deaths as contrasts of interest. Using psychiatric comorbidity (one of two data reduction identified factors, see methods) as the contrast of interest resulted in 49 DEGs (adjusted q-value <=0.05) in sgACC and 3 DEGs in Ant-Ins. Suicide completion, our mortality outcome of interest, was associated with 54 DEGs in sgACC and 6 DEGs in Ant-Ins. Longevity (i.e., another data reduction identified a factor measuring higher age at death and related variables despite chronic lifetime psychiatric and medical illnesses), on the other hand, was associated with more DEGs in Ant-Ins (145 DEGs) than in sgACC (14 DEGs).

### Suicide completion-related differential gene expression analysis identified DEGs

Because a significant percentage of our studied mood disorder samples died by suicide/are suicide completers (∼60+% of the included mood disorder donor samples), this makes suicide a significant predictor of premature death and underlying DEGs in our disease samples. We, therefore, quantified DEGs for suicide completion vs. non-suicide deaths (excluding controls which, by default, had no mental disorder or suicide history) at p<=0.05 FDR. We found <u>six</u> downregulated Ant-Ins DEGs, including the cell growth inhibiting serpine family *SERPINA3* gene that was earlier implicated in schizophrenia, *FOSB* transcription factor involved in encoding leucine zipper proteins and dimerization of proteins of the JUN family, thereby regulating leukocyte and T-cell proliferation, differentiation, and transformation (Heximer et al. 1996; Baumann et al. 2003; Nestler 2015; Manning et al. 2017), inflammation and tissue remodeling *CHI3L1* gene found to be associated with Alzheimer’s disease and schizophrenia (Su et al. 2022), and a BAALC-AS1 lncRNA (Punzi et al. 2019) (**Table S1A, Fig 4A-B**), as well as a long non-coding RNA AC145676.2, etc. The GO-terms for Ant-Ins DEGs in suicide completion were enriched for tyrosine-protein processing, B-cell activation, immune responsiveness, and regulatory T-cell pathways (Kathryn et al. 2018) (**Fig 4A-B)**.

**Figure 4.**
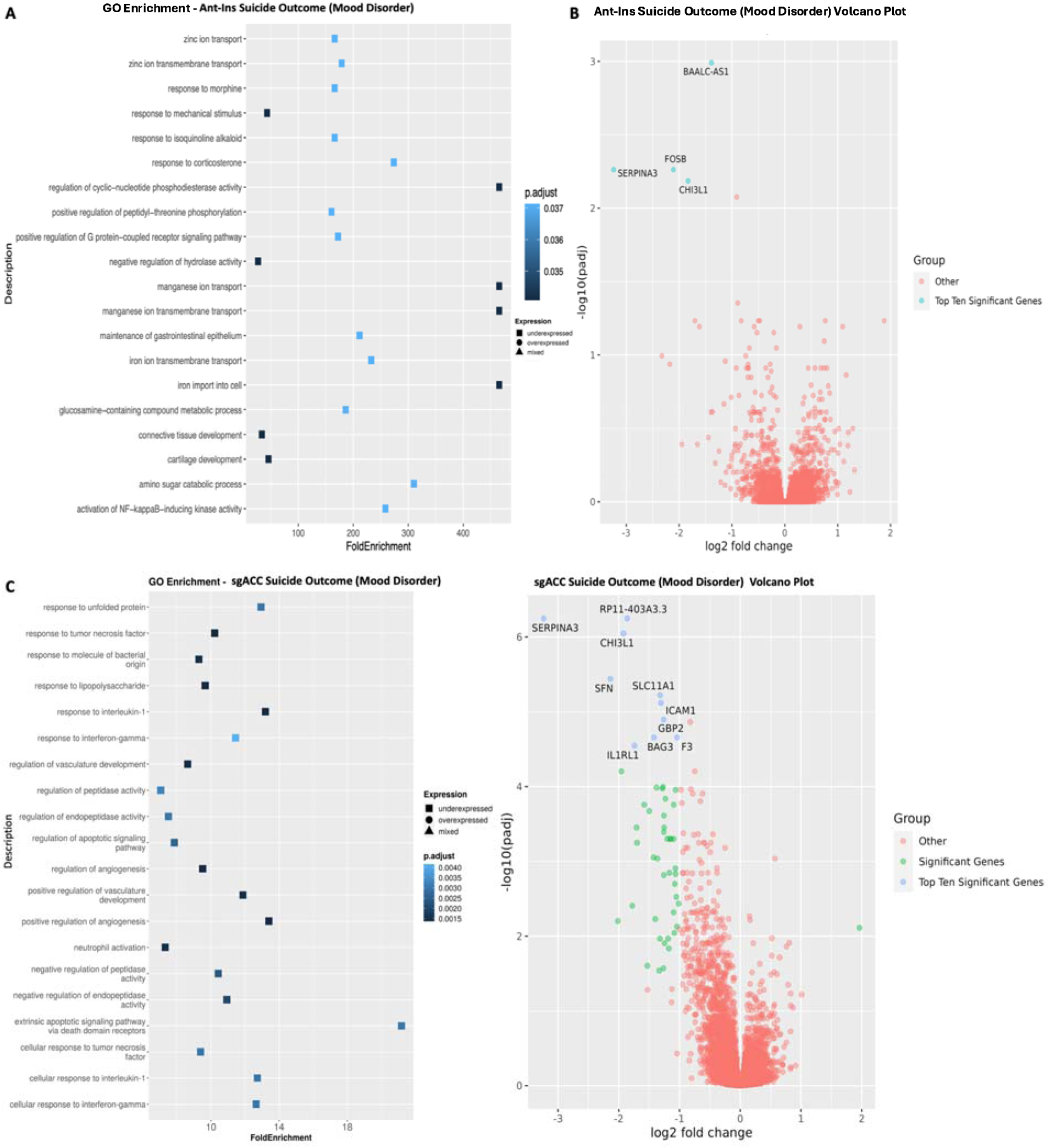
Gene Ontology (GO) terms and Volcano plots for Suicide Completions vs. Non-Suicide Deaths. **A**) illustrates GO terms for GECs in Ant-Ins for suicide completion vs. non-suicide deaths, with **B**) depicting the related volcano plot for the Ant-Ins results in **A**. **C & D**) illustrates GO terms in sgACC representing suicide completion vs. non-suicide deaths, and related volcano plot for the sgACC results in **C**. Because the -log10 (adjusted q-values) of a large number of genes in Insula was close to zero, volcano plots were generated using q-value instead of adjusted q-value. Genes meeting the following cutoffs: adjusted q-value <0.05 and absolute log2 fold change >= 1 were highlighted on the volcano plot as significant genes.

Our analysis of DEGs in the sgACC associated with suicide completion vs. non-suicide deaths (excluding controls) identified predominantly downregulated DEGs, including downregulated markers (i.e., 313 of 332 markers were downregulated), including <u>three</u> members of the cell growth inhibiting serpine family of proteins *SERPINA3, SERPINE1*, and *SERPINA1* (implicated in schizophrenia, Alzheimer’s, and Parkinson’s diseases and Ant-Ins regional DEGs in suicide completers), the inflammation mediator *CHI3L1* (downregulated in Ant-Ins in suicide completers). Furthermore, the interleukin *IL1RL1* (Kathryn et al. 2018) and *HILPDA* autocrine signaling lipid storage genes (Cantarelli et al. 2014) were also downregulated, alongside an additional 307 downregulated DEGs in the sgACC of suicide completers. Conversely, the N-acetyltransferase meta-pathway biotransformation I and II gene, *NAA40*, the helix-loop transcriptional regulator *NPAS4*, and 17 other DEGs were selectively upregulated in sgACC of suicide death cases (**Table S1B; Fig 4C-D**). GO-terms for suicide completion-associated DEGs in sgACC were enriched for innate and adaptive immune/inflammatory, cell-type mediated tissue remodeling, and apoptosis regulatory genes (Raja et al. 2022) (**Fig 4C-D)**. For results of overlaps between the Ant-Ins and sgACC, see our Rank Rank Hypergeometric findings (*RRHO,* **Fig 5**).

**Figure 5.**
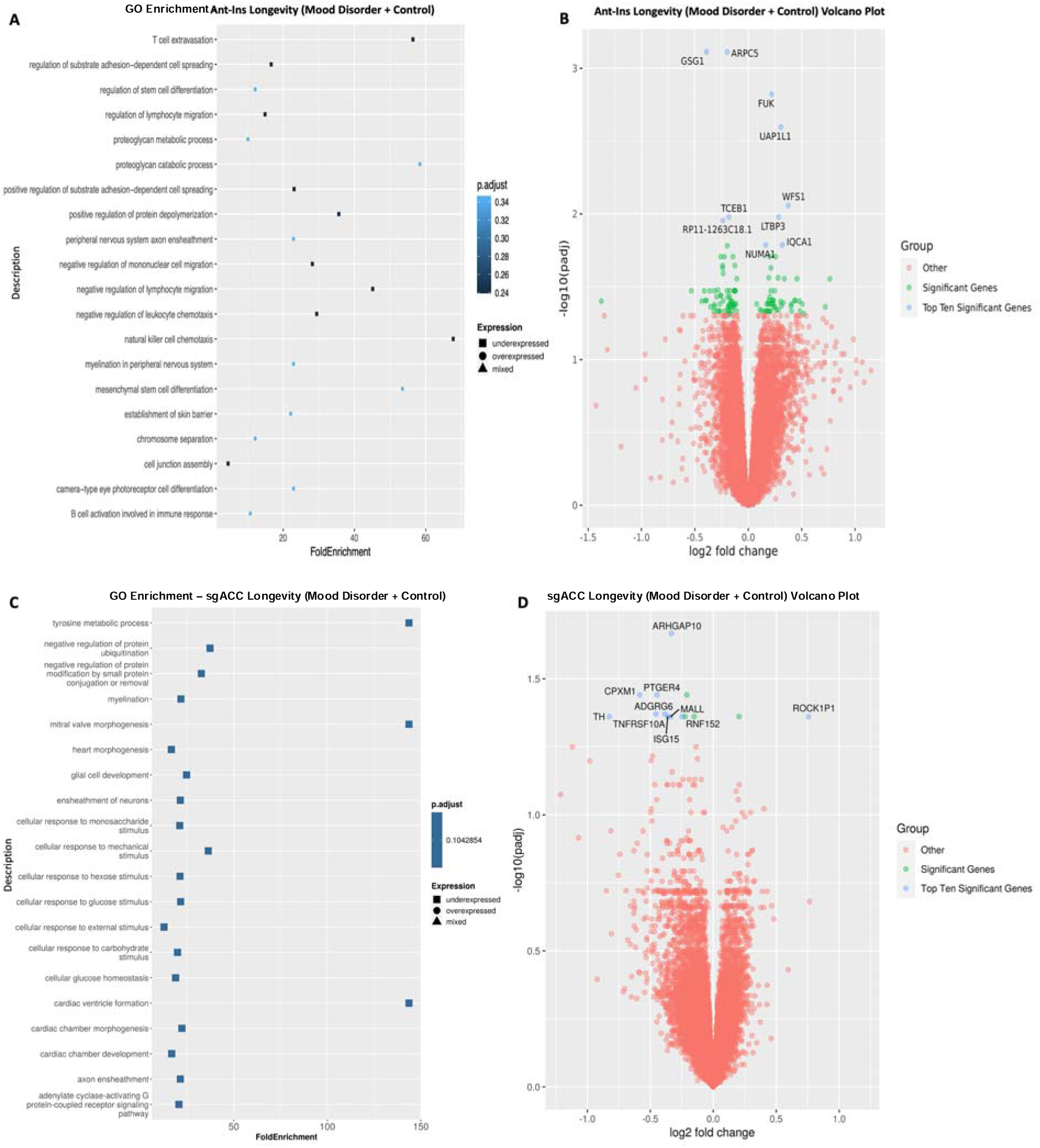
Gene Ontology (GO) terms and Volcano plots for High vs Low Longevity (higher-order factor). **A**) illustrates GO terms for GECs in Ant-Ins for high longevity vs. low longevity in all samples, with **B**) depicting the related volcano plot for the Ant-Ins results in **A** for the combination of mood disorders and unaffected control samples. **C & D**) illustrates GO terms in sgACC representing high longevity vs. low longevity for all samples, and related volcano plot for the sgACC results in **C**. Because the -log10 (adjusted q-values) of a large number of genes in Insula was close to zero, volcano plots were generated using q-value instead of adjusted q-value. Genes meeting the following cutoffs: adjusted q-value <0.05 and absolute log2 fold change >= 1 were highlighted on the volcano plot as significant genes.

**Figure 6.**
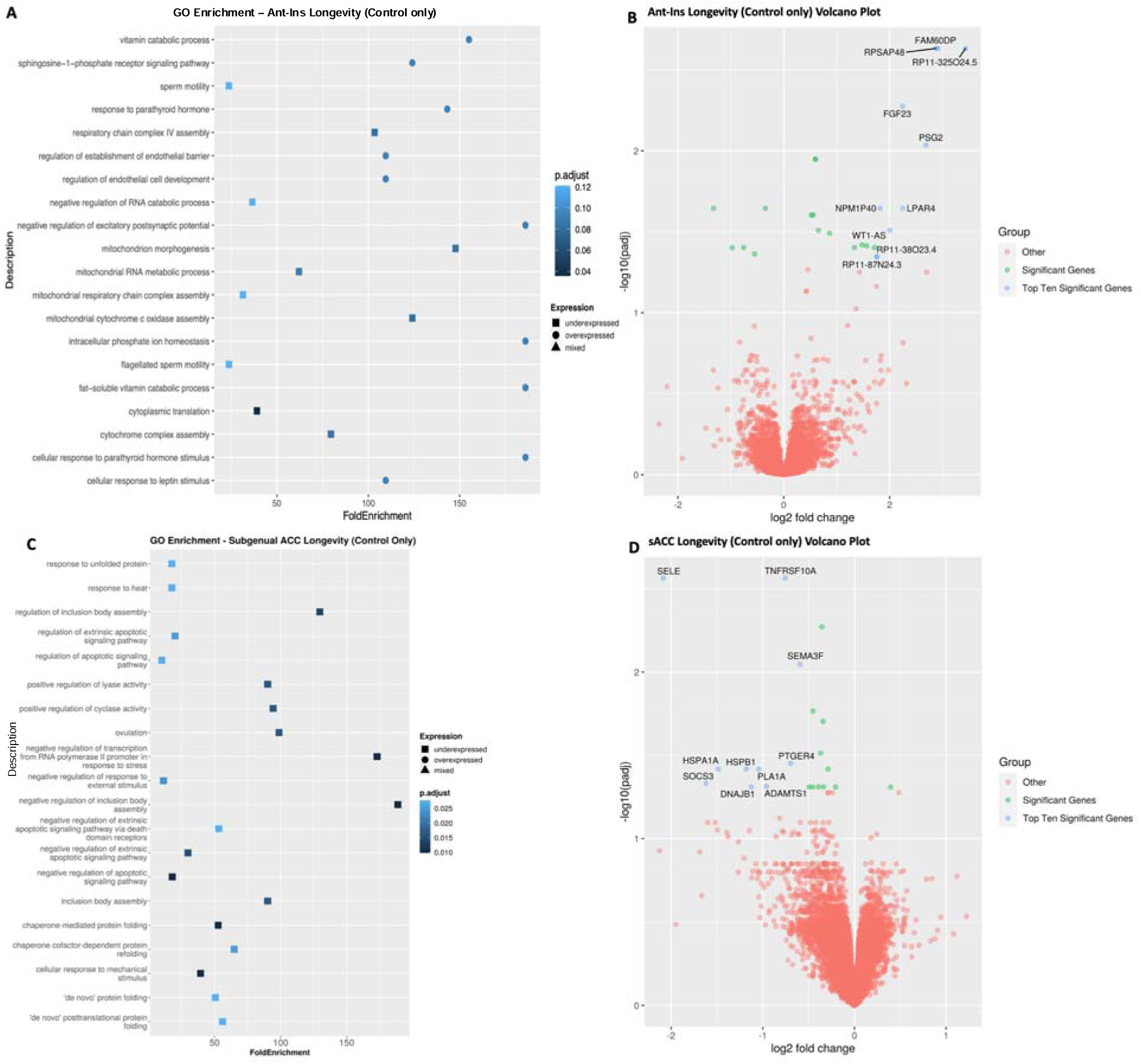
Gene Ontology (GO) terms and Volcano plots for High vs Low adaptive Longevity (in controls only). **A**) illustrates GO terms for GECs in Ant-Ins for high longevity vs. low longevity in controls, and **B**) related volcano plot for the results in **A** for unaffected controls. **C & D**) illustrates GO terms in sgACC for high adaptive longevity vs. low adaptive longevity in unaffected controls, and related volcano plot in **C**. Because the -log10 (adjusted q-values) of a large number of genes in Insula was close to zero, volcano plots were generated using q-value instead of adjusted q-value. Genes meeting the following cutoffs: adjusted q-value <0.05 and absolute log2 fold change >= 1 were highlighted on the volcano plot as significant genes.

### Longevity-related differential gene expression analysis identified DEGs

#### Longevity-associated DEGs in mood disorders and unaffected controls

Next, we assessed longevity-associated DEGs across all samples for the Ant-Ins and sgACC separately. We applied an identical median split-half comparison as in our *psychiatric morbidity* analysis to identify DEGs associated with high vs. low *longevity* at adjusted p<=0.05 FDR. We found that high vs. low *longevity* was associated with <u>eighty-two</u> downregulated Ant-Ins DEGs, including the protein synthesis *PSK5* (downregulated in Ant-Ins high *psychiatric morbidity* associated DEGs), cellular actin polymerization *ARPC5*, and RNA polymerase binding *GSG1* genes (**Table S2A; Fig 5A-B**). Conversely, we found <u>sixty-four</u> upregulated genes, including the glycoprotein and glycolipid synthesizer and carbohydrate metabolizer *FUK* (Cantarelli et al. 2014), UDP-N-acetylglucosamine biosynthetic processor *UAP1L1*, and cellular calcium regulator wolframin *WFS1* genes (Koido et al. 2005; Kato et al. 2008; Munshani et al. 2011; Seifuddin et al. 2013; Xavier et al. 2016) in the Ant-Ins (**Table S2A; Fig 5C-D**). Together, the GO-term pathways for *longevity-*associated Ant-Ins DEGs were enriched for protein synthesis, synaptic membrane, and receptor signaling genes (Manji & Chen, 2000; Laguesse & Ron 2020; Szczepankiewicz et al. 2021; Akula et al. 2021) (**Fig 5**). Similar analysis of high vs. low *longevity* (including MDD, BD, and controls) in the sgACC yielded <u>no</u> DEGs at adjusted q-values of q<=0.01 for high vs. low *longevity* in the combined mood disorders and the unaffected controls.

#### Longevity-associated DEGs in mood disorders

We compared high vs. low *longevity* exclusively in the mood disorder samples (excluding controls) to assess DEGs for maladaptive aging, first in the Ant-Ins followed with the sgACC analysis at p<=0.05 FDR. We found <u>one hundred and</u> <u>twenty-nine</u> downregulated DEGs in the Ant-Ins, including the protein synthesis *PSK5* gene and MTCO2P1*2* pseudogene, and <u>sixty-four</u> upregulated genes (**Table S2A)**. Unlike the lack of DEGs in the sgACC of the combined samples of mood disorders and control, our assessment of DEGs associated with *longevity* in mood disorders only (excluding controls) yielded <u>ten</u> downregulated genes, including GTPase activator and cellular cytoskeletal and apoptosis regulator *ARHGAP10* gene previously implicated in brain morphogenesis and schizophrenia (Kim et al. 2003; Sekiguchi et al. 2020; Hada et al. 2021), and <u>twelve</u> upregulated genes, including protein-kinase ROCK1P1 pseudogene (**Table S2B**). The GO-terms for *longevity*-associated DEGs in sgACC identified enriched pathways for tyrosine and protein synthesis and protein folding, cellular apoptosis, body assembly, and negative transcriptional regulatory genes (Jope 1999; Alpert & Fiori 2014; Cox et al. 2021) in mood disorders only cohort.

#### Longevity-associated DEGs validated in unaffected controls

Unlike *psychiatric morbidity*, the presence of which was an exclusion criterion for controls, the phenotypic loading for our identified higher-order *longevity* factor (which reflects variability in *a.* marital status, *b.* # of children, *c.* Axis-III comorbidity, and *d.* age at death) was naturally expected to be more normally distributed across all samples including controls. As such, we treated variability in *longevity* in controls as a proxy measure of how socially enriched the donors’ lives were (marital status & #of children) in addition to how long they lived despite the presence of chronic medical diseases (Axis-III morbidity and age at death). With the adaptive measure of *longevity* in controls in mind, we conducted a differential gene expression analysis of high vs. low *longevity* exclusively in the unaffected controls using the p<=0.05 FDR threshold to identify DEGs associated with mentally adaptive/resilient *longevity* (i.e., having no recorded lifetime history of *psychiatric morbidity*). Comparing high vs. low adaptive *longevity* in controls only, we found <u>thirty-four</u> upregulated DEGs in the Ant-Ins (**Table S3A**), including the GTPase and metal iron binding gene implicated in autism AGAP7P gene, G-protein coupled receptor S1PR2, and LPAR4 genes, Leucine-rich LRRC69 gene, DNA binding transcription factor SHOX, antisense RNA WT1-AS gene, fibroblast growth factor 23 *FGF23* anti-aging gene (Haussler et al. 2016), the pregnancy-specific Beta-1-Glycoprotein 2 *PSG2* (Khan et al. 1992) gene, and several other genes and pseudogenes (**Table S3A; Fig 6A**). We further found <u>twenty-five</u> downregulated DEGs (**Table S3A**) associated with adaptive *longevity* in controls only in Ant-Ins, including AC007192.6/PIK3R2 gene involved in neurodevelopment, and the collagen type VI alpha chain COL6A3 gene associated with connective tissue/muscle regeneration and disease (**Table S4A**), etc. Assessing the GO-terms for *longevity*-associated DEGs in Ant-Ins of controls revealed pathways enriched for cellular response to vitamin D metabolic processes, lipid metabolism, and cellular homeostasis (Cantarelli et al. 2014; Vandekopple et al. 2019) (**Fig 6A-B)**.

High vs. low adaptive *longevity*-associated DEGs in the sgACC of controls only identified o<u>ne</u> upregulated uncharacterized marker STX16-NPEPL1. However, high longevity versus low longevity controls comparison in the sgACC resulted in <u>twenty</u> downregulated genes, including the blood leukocyte chaperoned cytokine-stimulated *SELE*; and tumor necrosis factor-related apoptosis inducer *TNFRSF10A* (Uversky 2014; Miller & Raison 2016; Kathryn et al. 2018), as well as *TMEM45B*, *SEMA3F*, *ADGRL4*, *VASP*, *SOCS3*, *ADAMTS1*, *DNAJB1*, *ICAM2*, and *NOS3* genes. Further downregulated DEGs associated with adaptive *longevity* in sgACC include the major histocompatibility complex-heat shock protein *HSPA1A* and *HSPB1*, as well as *PLA1A* and *CNN2* genes, and an uncharacterized PUDP, AF131216.6 genes (**Table S3B**). Additionally, these downregulated DEGs included the calcium membrane *ORAI1* that channels calcium influx into T-Cells (Luik & Lewis 2007; Miller 2010; Vaeth et al. 2020; Ramesh et al. 2021; Voros et al. 2021), the T-cell-signaling G-protein-coupled prostaglandin E receptor *PTGER4* gene (Hsiao et al. 2021), and the synaptic membrane/GTPace signal transducers *PLEKHG1* genes (**Table S3B; Fig 6C-D**). GO-terms for adaptive *longevity*-associated DEGs in sgACC were enriched for tyrosine or protein synthesis/folding, apoptosis, body assembly, and negative transcriptional regulation (Jope 1999; Alpert & Fiori 2014; Rangaraju et al. 2016; Cox et al. 2021) (**Fig 6C-D)**.

***Ant-Ins and sgACC Rank Rank Hypergeometric Overlap (RRHO) in Psychiatric Morbidity, Suicide Completion, and Longevity:*** In line with previous neuroimaging findings of similar patterns of reduced anatomical gray matter reductions in neuropsychiatric diagnoses, suicidal phenotypes, and aging, we found a preponderance of gene expression overlap across the AIAC network. Specifically, we found both downregulated and upregulated gene expression overlaps across the AIAC network such that all three contrasts of interest showed extensive interregional overlap. However, the degree of overlapping downregulated gene expression was highest in suicide completion and longevity associated DEGs, respectively (**Fig 7A-C**).

**Figure 7.**
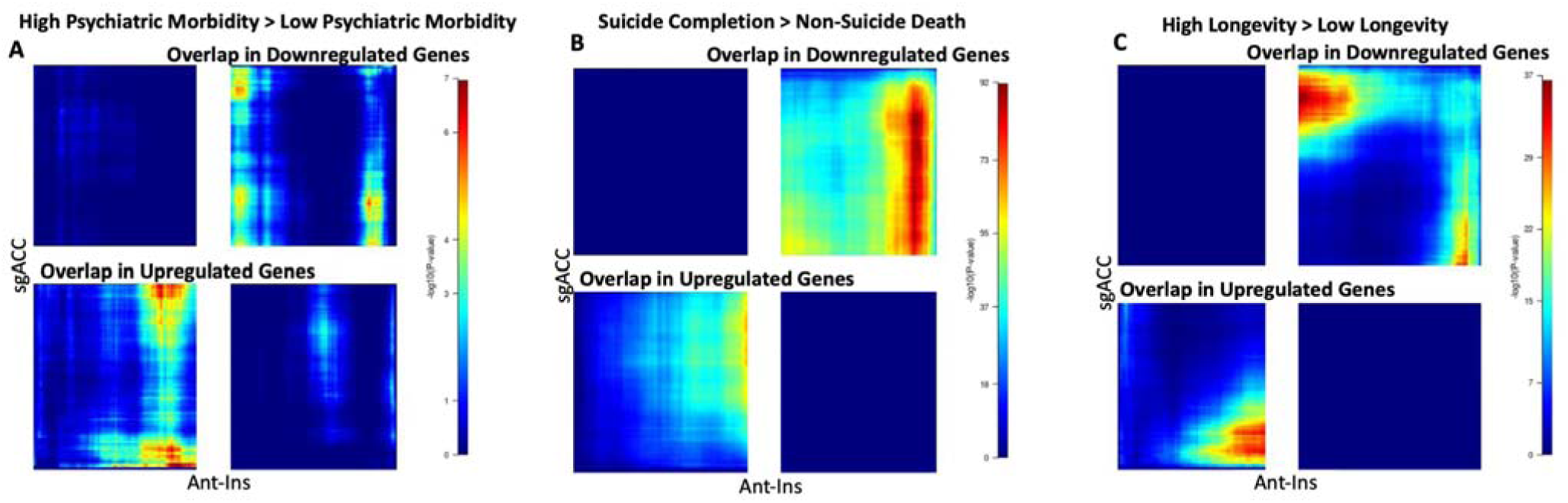
RRHO Enrichment Results. The significant heatmaps illustrates quadrants (top right) or (bottom left) of hypergeometric overlap in gene expression changes in the same direction in Ant-Ins and sgACC regions of a AIAC network. **A**) Shows that for high versus low psychiatric morbidity, there are inter-regional overlaps in both downregulated and upregulated gene expression. B) For suicide completion however, the preponderance of overlap in Ant-Ins and sgACC regional gene expression was predominantly downregulated, similar to longevity associated gene expression patterns observed in C. The color bars on the right represents the values of the log10 p-values of the correlations.

## DISCUSSION

This study identified patterns of differentially expressed genes (DEGs) underlying mood disorders and co-occurring psychiatric disorders. We applied novel data reduction techniques to study the molecular neurobiology of a critical AIAC brain network. We demonstrated that it is feasible to identify specific DEGs for complex disease phenotypes like *psychiatric morbidity* and suicide outcomes (which are composite measures of maladaptive mood and related behavioral outcomes) in addition to *longevity* (both longevity while having a lifelong mental illness and longevity in the absence of mental illness). Using a postmortem brain RNA-sequencing approach, we focused on the AIAC brain network, known to be involved in the regulation of affect and mood in health and disease (Nauta 1971; Goldman-Rakic 1988; Drevets et al. 2008), and known to harbor reduced anatomical integrity in individuals with mood and comorbid psychiatric disorders (Savitz & Drevets 2009; 3 et al. 2015; Wise et al. 2017; Jabbi et al. 2020a) and suicidal phenotypes (Schmaal et al. 2020a; Jabbi et al. 2020b). The targeted AIAC network exhibits anatomical and physiological changes associated with therapeutic responses (Salvadore et al. 2009; McGrath et al. 2013; Riva-Posse et al. 2018), and transcriptome studies of this brain network revealed molecular abnormalities associated with mood disorders and suicide phenotypes (Galvalvy et al. 2013; Orozco-Solis et al. 2017; Lutz et al. 2017; Zhou et al. 2018).

To study a more specific AIAC network molecular repertoire underlying mood disorders and related *psychiatric morbidity* and mortality, we used WGCNA to assess global gene co-expression related to mood disorder phenotypes. We found no specific coexpressed networks associated with the diagnoses of MDD or BD in the AIAC network. In contrast to the traditional diagnostic focus of identifying gene expression correlates for MDD and BD vs. controls, which in this study were not correlated with any significant WGCNA modules, our WGCNA analysis revealed significant negative correlations between the higher order factors such as *psychiatric morbidity*, *longevity/aging* and related suicide variables recapitulates previous findings by revealing enriched co-expression modules related to critical cellular processes like synaptic membrane and ion channel signaling (Sokolowski et al. 2018; Pishva et al., 2017; Akula et al. 2021; Zandi et al. 2022), and critical homeostatic regulatory processes such as inflammatory signaling (Elhaik & Zandi 2015; Shipweck et al. 2020; Arasappan et al. 2021), metabolic processes/mitochondrial translation (Laguesse & Ron 2020), and ATP-synthesis pathways (Traub & Lichtstein 2000; Innes et al. 2019) in the Ant-Ins for *psychiatric morbidity*. In line with our WGCNA findings in the Ant-Ins, the co-expression modules in the sgACC revealed correlations between *psychiatric morbidity* and *longevity/aging* with enriched protein synthesis (Pishva et al., 2017; Akula et al. 2021), neurodegeneration, basic cellular processes and inflammatory signaling (Glover et al., 2018; Raja et al. 2022). Together, these results underscore the complexity of AIAC network’s molecular processes in health and disease across the human lifespan, and reveals a molecular dysregulations that are likely proximate to homeostatic imbalances associated with dysregulated mood states.

As humans develop from birth to adulthood and gradually attain advanced aging, repeated exposures to social and environmental pathogens can negatively impact brain and body functions, leading to various maladaptive phenotypes and diseases like mood disorders. It is, therefore, plausible that the phenotypic expression of lifetime resilience to mood disorders and co-occurring chronic medical diseases is essential for the long-term maintenance of well-being and longevity. While previous studies have examined the relationship between lifetime mood disorder risks and longevity, using different metrics such as years of life lost, see an influential review on this (Murray et al. 2012), the neurobiology underlying variability in mood disorder morbidity and related mortality phenotypes remains obscure. This study is the first to systematically characterize the critical brain molecular mechanisms with dosage-sensitive relationships in mood disorders and related mortality/suicide outcomes by highlighting the downregulatory vs. upregulatory gene expression repertoires associated with complex morbidity and mortality phenotypes in mood disorders. Although few studies (Turecki et al. 2019), including our recent work (Jabbi et al. 2020a; Akula et al. 2021; Arasappan et al. 2021) have characterized maladaptive (lifetime morbidity and suicide mortality) relative to adaptive (longevity despite lifetime psychiatric and other diseases) phenotypes in postmortem brain network transcriptome studies of mood disorders, our current direct comparisons between MDD vs. controls and BD vs. controls identified fewer mood disorder diagnoses-specific DEGs relative to our comparisons of higher-order factors such as high vs. low *psychiatric morbidity* and high vs. low *longevity* across the AIAC network. Notably, even after excluding controls to assess if our identified DEGs are more proximate to mood pathology, our observation of downregulated protein synthesis was preserved, suggesting a more perversive protein synthesis dysregulation at the level of DEGs in increased *psychiatric morbidity* than previously thought. Further analysis of high vs. low *psychiatric morbidity* in the sgACC yielded downregulated G-protein-coupled DEGs, as well as a predominant upregulation of metabolic, stress-responsive (Cantarelli et al. 2014; Vandekopple et al. 2019), inflammatory, and synaptic membrane/calcium ion channel (Sokolowski et al. 2018), and iron homeostasis DEGs.

By assessing DEGs for suicide completion, we uncovered predominantly downregulated Ant-Ins DEGs associated with protein synthesis, immune and inflammatory signaling, B-cell activation (Miller & Raison 2016), and synaptic membrane regulatory functions (Sokolowski et al. 2018; Zandi et al. 2022). In line with the Ant-Ins findings, sgACC DEGs associated with suicide completion were predominantly downregulated genes, including innate and adaptive immune/inflammatory pathways (Uversky 2014; Mechawar & Savitz 2016; Kathryn et al. 2018; Giridharan et al. 2020), and tissue/cellular development and apoptosis regulatory pathways, as well as translational regulatory pathways. Given that increased *psychiatric morbidity* is the most predominant risk factor for suicide, these findings can be interpreted in two ways. *First*, Adverse life experiences and stress are likely causally linked to mood disorders and related reductions in AIAC brain network, as well as with increased brain synaptic imbalances that can trigger microglial mediated overprunning of stress-damaged neurons and result in loss of neuronal connectivity and related synaptic membrane gene regulatory functions, with potential developmental and lifelong impaired functional consequences (Glover et al., 2018; Raja et al. 2022). *Second*, our findings of neurodevelopmental, tissue developmental, and synaptic/ion channel regulatory DEGs can be meaningful both in terms of early developmental gene-mediated neurodevelopmental deficits leading to lifelong repercussions such as mood regulatory vulnerabilities and increased *psychiatric morbidity* and suicide risk outcomes (Turecki et al. 2019; Jabbi et al. 2020a; Punzi et al. 2022). Most importantly, cellular and neurodevelopmental gene regulatory abnormalities may comprise a neurobiological vulnerability that manifests as a mood disorder when individuals with these developmental gene-regulation deficits are further exposed to early life adversities, as is often the case for individuals with high familial/genetic risk for mood disorders (Slavich et al. 2010; Cantarelli et al. 2014; Cox et al. 2021).

In line with increased risk for adverse childhood traumatic experiences coupled up increased likelihood of socioeconomic adversity for individuals with familial risk for mood disorders, our postmortem mood disorder samples comprising 112 of the 180 donors, we found that the observed DEGs in the Ant-Ins network in relation to high vs. low *longevity* in controls were associated with downregulation in protein synthesis and upregulated synaptic membrane signaling. On the one hand, our observed *longevity*-associated DEGs specific to the controls, akin to adaptive *longevity/aging*, were both upregulated in biological systems associated with enhancing anti-aging or longevity processes, as well as in cellular homeostatic processes such as response to vitamin D metabolic processes and lipid metabolism (Cantarelli et al. 2014; Vandekopple et al. 2019). Adaptive *longevity*/no lifetime psychiatric disease in controls was additionally associated with downregulated DEGs related to neurodevelopment, and connective tissue and muscular degeneration, potentially reflecting the aging and physical disease-related brain molecular correlates for resilience to mood dysfunctions. On the other hand, DEGs for high vs. low adaptive *longevity* in sgACC in controls were associated with downregulated cellular apoptosis, negative regulation of protein synthesis and folding, negative regulation of stress-induced transcriptional activity, and response to heat shock. The findings of downregulated apoptotic and inhibitory processes for adaptive biological functions in individuals who died without ever having any know psychiatric disorders suggest a proximate role for the sgACC cellular integrity in maintaining functional behavioral health across the lifespan. Future studies will be needed to validate these region-specific findings. However, our results of adaptive *longevity*-associated upregulatory anti-aging molecular pathway functions, coupled with pregnancy-specific, and cellular metabolic/homeostatic DEGs in Ant-Ins, as opposed to downregulatory DEGs related to accelerated cell death and inhibitory processes for protein synthesis in sgACC in controls with no history of psychiatric diseases, strongly point to possible adaptive molecular processes that may be necessary for the maintenance of basic cellular and neuronal regulation of complex affective behaviors in the AIAC brain network. If replicated, these observed novel mechanisms may be critical to the basic molecular regulation of longevity, even in age-related comorbid medical conditions.

Here, we tested the hypothesis that the phenotypic complexity of comorbid psychiatric diseases in individuals with primary mood disorders may confer immune and inflammatory-related neuropathological markers that may also be involved in longevity/aging and related lifetime chronic physical disease. Our results generally supported this hypothesis and underscored the need for studying specific immune/inflammatory mechanisms for mood disorder phenotypes. Given that mood disorders and related psychiatric conditions are well documented to be triggered by stressful/traumatic environmental experiences which can cause a long-term cascade of immune and inflammatory processes related both physical and psychological harm resulting from the noxious environmental exposures, it is likely that the overlap in the downregulated immune and inflammatory responses related to both mental and physical diseases identified in our samples are evidence of shared pathobiological processes underlying immune clearance of disease induced inflammation that can originate from both psychological/environmental trauma and physical disease mechanisms. Furthermore, our findings of downregulated protein synthesis and ion/calcium channel signaling DEGs in high vs. low *psychiatric morbidity*, shown to be spread across the entire studied network, could serve as a possible underlying mechanism for the often-observed anatomical integrity reductions in this brain network in in mood disorders and comorbid conditions (Goodkind et al. 2015; Wise et al. 2017; Jabbi et al. 2020a).

The current study has limitations in that even though we downsampled the sgACC data to be comparable with the Ant-Ins data, the RNAseq methods were not identical across the two regions, so it we cannot preclude the possibility that some differences we observed between Antins and sgACC regions are driven by methodologic differences. However, such differences cannot explain the convergent signals we observed. Secondly, even though 75 of the 100 donors we studied contributed data from both Ant-Ins and sgACC regions, there were more than twice as many donors of sgACC tissue. This imbalance means that the sources of variability likely differed across the two brain regions, reducing comparability. Thirdly, bulk RNA sequencing cannot account for cell-type specific differences in gene expression signatures in terms of which cell-types are driving specific downregulation or upregulation of key transcriptional elements. Future studies that apply novel single cell approaches using the approaches defined in our study will be needed to identify dosage-sensitive cell-type specific neuropathological influences at transcriptomic scales and guide novel diagnostic and therapeutic advances. Finally, gene expression changes in post-mortem tissue may not capture changes over the lifespan and could result, in part, from illnesses or their treatment.

Our findings of an association between chronic *psychiatric morbidity* (higher numbers of lifetime *psychiatric disease comorbidity*, and *longevity/aging*), as well as completed suicide, with a preponderance of downregulated protein synthesis and immune/inflammatory, cellular developmental, and metabolic DEGs, likely underscores the important role of these molecular mechanisms in the maintenance of brain and body homeostasis in health and diseases. The morbidity and mortality-related gene expression changes we observed highlight key immune-metabolic and cellular signaling pathways within a critical, hardwired AIAC brain network involved in emotional and mood regulatory functions. In conclusion, our findings provide a mechanistic framework for understanding dosage-dependent (i.e., downregulated vs. upregulated) gene expression repertoires for adaptive and maladaptive mood functions, and could inform novel diagnostic and therapeutic innovations for comorbid psychiatric disease phenotypes, and suicide mortality outcomes across the lifespan.

## Acknowledgements

The NIMH Human Brain Collection Core (HBCC) provided RNA-samples for all 100 postmortem Anterior Insula. While the data for the 152 subgenual anterior cingulate brain donors was included under a data transfer agreement with the NIMH. We thank Drs. Stefano Marenco, Pavan Auluck and HBCC colleagues for providing the samples and related data. We thank Jessica Podnar and the University of Texas genome sequencing and facility (GSAF) colleagues for RNA-seq support. This work was supported by the Dell Medical School Mulva Clinics for the Neurosciences, UT Austin, and by 5R21MH115326-02 and 1R01MH134791-01 from NIMH to MJabbi. NAkula and FJMcMahon are supported by ZIAMH002843-20 and 1ZIAMH002843-20 from NIMH. Supported in part by the Intramural Research Program of the NIMH.

**Tables 1**-**6** Gene expression differences detected at adjusted p<0.05. Gray shaded results represent downregulated genes (negative Log2Foldchange values), whereas non-shaded results represent upregulated genes (positive Log2Foldchange values).

## Notes

### Competing Interest Statement

The authors have declared no competing interest.

## REFERENCES

Kessler, RC, et al. Prevalence, Severity, and Comorbidity of Twelve-month DSM-IV Disorders in the National Comorbidity Survey Replication (NCS-R). Archives of General Psychiatry. 2005; 62(6): 617–627.

Murray, CJL et al. Disability-adjusted life years (DALYs) for 291 diseases and injuries in 21 regions, 1990–2010: a systematic analysis for the Global Burden of Disease Study 2010. The Lancet; 2012; 380(9859): 2197–2223.

Oquendo MA., Currier D, Liu SM, Hasin D, Grant G, and Blanco C. Increased risk for suicidal behavior in comorbid bipolar disorder and alcohol use disorders: results from the National Epidemiologic Survey on Alcohol and Related Conditions (NESARC). The Journal of Clinical Psychiatry. 2010; 71(7): 902–9.

Whiteford HA, Degenhardt L, Rehm J, Baxter AJ, Ferrari AJ, Erskine HE, Charlson FJ, Norman RE, Flaxman AD, Johns N, Burstein R, Murray CJ, Vos T. Global burden of disease attributable to mental and substance use disorders: findings from the Global Burden of Disease Study 2010. Lancet. 2013; 382(9904): 1575-86.

Baxter AJ, Harris MG, Khatib Y, Brugha TS, Bien H, Bhui K. Reducing excess mortality due to chronic disease in people with severe mental illness: meta-review of health interventions. Br J Psychiatry. 2016; 208(4): 322–9.

Amare AT, Schubert KO, Klingler-Hoffmann M, Cohen-Woods S, Baune BT. The genetic overlap between mood disorders and cardiometabolic diseases: a systematic review of genome-wide and candidate gene studies. Transl Psychiatry. 2017; 7(1): e1007.

Niculescu AB, Le-Niculescu H, Levey D, Phalen P, and Dainton H. Precision medicine for suicidality: from universality to subtypes and personalization. Molecular Psychiatry. 2017; 22(9): 1250–1273.

Turecki G, Brent DA, Gunnell D, O’Connor RC, Oquendo MA, Pirkis J, Stanley BH. Suicide and suicide risk. Nat Rev Dis Primers. 2019; 5(1): 74.

Gibbs CR, Blann AD, Watson RD, Lip GY. Abnormalities of hemorheological, endothelial, and platelet function in patients with chronic heart failure in sinus rhythm: effects of angiotensin-converting enzyme inhibitor and beta-blocker therapy. Circulation. 2001; 103(13): 1746–51.

Dum RP, Levinthal DJ, Strick PL.The mind-body problem: Circuits that link the cerebral cortex to the adrenal medulla. Proc Natl Acad Sci U S A. 2019; 116(52): 26321–26328.

Levinthal DJ, Strick PL. Multiple areas of the cerebral cortex influence the stomach. Proc Natl Acad Sci U S A. 2020; 117(23): 13078–13083.

Jabbi M, Bastiaansen J, and Keysers C. A common anterior insula representation of disgust observation, experience, and imagination shows divergent functional connectivity pathways PLoS One. 2008; 3(8): e2939.

Craig, A. D. How do you feel - now? The anterior insula and human awareness. Nature Reviews Neuroscience. 2009; 10(1): 59–70. 1

Khalsa SS, Adolphs R, Cameron O, Critchley H, Davenport P, Feinstein J, Feusner J, Garfinkel S, Lane R, Mehling W et al. Interoception and Mental Health: A Roadmap. Biological Psychiatry: Cognitive Neuroscience and Neuroimaging. 2018; 3(6): 501–513.

Nauta, WJH. The problem of the frontal lobe: A reinterpretation. Journal of Psychiatric Research. 1972; 8(3): 167–87.

Goldman-Rakic PS. Topography of Cognition: Parallel Distributed Networks in Primate Association Cortex. Annual Review of Neuroscience. 1988; 11(1): 137–156.

Harrison NA, Brydon L, Walker C, Gray M, Steptoe A, and Critchley H. Inflammation causes mood changes through alterations in the subgenual cingulate activity and mesolimbic connectivity. Biological Psychiatry. 2009; 66(5): 407–414.

Joyce MKP, and Barbas H. Cortical Connections Position Primate Area 25 as a Keystone for Interoception, Emotion, and Memory. Journal of Neuroscience. 2018; 38(7): 1677–1698.

Wang J, John Y, Barbas H. Pathways for Contextual Memory: The Primate Hippocampal Pathway to Anterior Cingulate Cortex. Cereb Cortex. 2021; 31(3): 1807–1826

Drevets WC, Savitz J, Trimble M. The subgenual anterior cingulate cortex in mood disorders. CNS Spectr. 2008; 13(8): 663–81.

Goodkind M, Eickhoff S, Oathes D, Jian Y, Chang A, Jones-Hagata L, Ortega B, Zaiko Y, Roach E, Korgaonkar M et al. Identification of a common neurobiological substrate for mental illness. JAMA Psychiatry. 2015; 72(4): 305–315.

Wise T, J. Radua E, Cardoner VN, Abe O, Adams T, Amico F, Cheng Y, Cole J, de Azevedo CMP et al. Common and distinct patterns of grey-matter volume alteration in major depression and bipolar disorder: evidence from a voxel-based meta-analysis. Molecular Psychiatry. 2017; 22 (10): 1455–1463.

Jabbi M, Arasappan D, Eickhoff S, Strakowski S, Nemeroff C, and Hofmann H. Neuro-transcriptomic signatures for mood disorders morbidity and suicide mortality. Journal of Psychiatric Research. 2020a; 127: 62–74.

Lipska BK, Deep-Soboslay A, Weickert CS, Hyde TM, Martin CE, Herman MM, Kleinman JE. Critical factors in gene expression in postmortem human brain: Focus on studies in schizophrenia. Biol Psychiatry. 2006; 60(6):650–8.

Akula N, Marenco S, Johnson K, Feng N, Zhu K, Schulmann A, Corona W, Jiang X, Cross J, England B, Nathan A, Detera-Wadleigh S, Xu Q, Auluck PK, An K, Kramer R, Apud J, Harris BT, Harker Rhodes C, Lipska BK, McMahon FJ. Deep transcriptome sequencing of subgenual anterior cingulate cortex reveals cross-diagnostic and diagnosis-specific RNA expression changes in major psychiatric disorders. Neuropsychopharmacology. 2021; 46(7): 1364–1372.

Andrews, S. (2010). FastQC:A Quality Control Tool for High Throughput Sequence Data [Online]. Available online at: http://www.bioinformatics.babraham.ac.uk/projects/fastqc/

Bray N. L et al. (2016). Near-optimal probabilistic RNA-seq quantification. Nat Biotechnol. 2016; 34(5):525-7.

Pertea M, Kim D, Pertea GM, Leek JT, Salzberg SL. Transcript-level expression analysis of RNA-seq experiments with HISAT, StringTie and Ballgown. Nat Protoc. 2016; 11:1650–67.

Langfelder P, Horvath S. WGCNA: an R package for weighted correlation network analysis. BMC Bioinformatics. 2008; 9:559.

Costello, A. B., & Osborne, J. (2005). Best Practices in Exploratory Factor Analysis: Four Recommendations for Getting the Most from Your Analysis. Practical Assessment Research & Evaluation, 10, 1–9.

Kuleshov MV, Jones MR, Rouillard AD, Fernandez NF, Duan Q, Wang Z, Koplev S, Jenkins SL, Jagodnik KM, Lachmann A, McDermott MG, Monteiro CD, Gundersen GW, Ma’ayan A. Enrichr: a comprehensive gene set enrichment analysis web server 2016 update. Nucleic Acids Res. 2016; 44(W1): W90–7.

Cahill KM, Huo Z, Tseng GC, Logan RW, Seney ML. Improved identification of concordant and discordant gene expression signatures using an updated rank-rank hypergeometric overlap approach. Sci Rep. 2018; 8(1):9588.

Schloss P, Henn FA. New insights into the mechanisms of antidepressant therapy. Pharmacol Ther. 2004; 102(1): 47–60.

Ryding E, Ahnlide JA, Lindström M, Rosén I, Träskman-Bendz L. Regional brain serotonin and dopamine transporter binding capacity in suicide attempters relate to impulsiveness and mental energy. Psychiatry Res. 2006; 148(2-3): 195–203.

Miller AH, Raison CL. The role of inflammation in depression: from evolutionary imperative to modern treatment target. Nat Rev Immunol. 2016; 16(1): 22–34.

Pishva E, Rutten BPF, van den Hove D. DNA Methylation in Major Depressive Disorder. Adv Exp Med Biol. 2017; 978: 185–196.

Benjamini Y and Hochberg Y: Controlling the false discovery rate: A practical and powerful approach to multiple testing. J Roy Stat Soc B Met. 57:289–300. 1995.

Uversky VN. Wrecked regulation of intrinsically disordered proteins in diseases: pathogenicity of deregulated regulators. Front Mol Biosci. 2014; 1: 6.

Arasappan D, Eickhoff SB, Nemeroff CB, Hofmann HA, Jabbi M. Transcription Factor Motifs Associated with Anterior Insula Gene Expression Underlying Mood Disorder Phenotypes. Mol Neurobiol. 2021; 58(5): 1978–1989.

Pantazatos SP, Huang YY, Rosoklija GB, Dwork AJ, Arango V, Mann JJ. Whole-transcriptome brain expression and exon-usage profiling in major depression and suicide: evidence for altered glial, endothelial and ATPase activity. Mol Psychiatry. 2017; 22(5): 760–773.

Laguesse S, Ron D. Protein Translation and Psychiatric Disorders. Neuroscientist. 2020; 26(1): 21–42.

Cantarelli Mda G, Tramontina AC, Leite MC, Gonçalves CA. Potential neurochemical links between cholesterol and suicidal behavior. Psychiatry Res. 2014; 220(3): 745–51.

VandeKopple MJ, Wu J, Baer LA, Bal NC, Maurya SK, Kalyanasundaram A, Periasamy M, Stanford KI, Giaccia AJ, Denko NC, Papandreou I. Stress-responsive HILPDA is necessary for thermoregulation during fasting. J Endocrinol. 2017; 235(1): 27–38.

Sokolowski M, Wasserman J, Wasserman D. Gene-level associations in suicide attempter families show an overrepresentation of synaptic genes and genes differentially expressed in brain development. Am J Med Genet B Neuropsychiatr Genet. 2018; 177(8): 774–784.

Zandi PP, Jaffe AE, Goes FS, Burke EE, Collado-Torres L, Huuki-Myers L, Seyedian A, Lin Y, Seifuddin F, Pirooznia M, Ross CA, Kleinman JE, Weinberger DR, Hyde TM. Amygdala and anterior cingulate transcriptomes from individuals with bipolar disorder reveal downregulated neuroimmune and synaptic pathways. Nat Neurosci. 2022; 25(3): 381–389.

Zhang HL, Wang XC, Liu R. Zinc in Regulating Protein Kinases and Phosphatases in Neurodegenerative Diseases. Biomolecules. 2022; 12(6): 785.

Heximer SP, Cristillo AD, Russell L, Forsdyke D R. Sequence analysis and expression in cultured lymphocytes of the human FOSB gene (G0S3). DRDNA and cell biology, 1996; 15(12): 1025–38.

Baumann S, Hess J, Eichhorst ST, Krueger A, Angel P, Krammer PH, Kirchhoff S. An unexpected role for FosB in activation-induced cell death of T cells. Oncogene, 2003; 22(9): 1333–9.

Punzi G, Ursini G, Viscanti G, Radulescu E, Shin JH, Quarto T, Catanesi R, Blasi G, Jaffe AE, Deep-Soboslay A, Hyde TM, Kleinman JE, Bertolino A, Weinberger DR. Association of a Noncoding RNA Postmortem With Suicide by Violent Means and In Vivo With Aggressive Phenotypes. Biol Psychiatry. 2019; 85(5): 417–424.

Raja GL, Subhashree KD, Kantayya KE. In utero exposure to endocrine disruptors and developmental neurotoxicity: Implications for behavioural and neurological disorders in adult life. Environ Res. 2022; 203: 111829.

Koido K, Kõks S, Nikopensius T, Maron E, Altmäe S, Heinaste E, Vabrit K, Tammekivi V, Hallast P, Kurg A, Shlik J, Vasar V, Metspalu A, Vasar E. Polymorphisms in wolframin (WFS1) gene are possibly related to increased risk for mood disorders. Int J Neuropsychopharmacol. 2005; 8(2): 235–44.

Kato T, Ishiwata M, Yamada K, Kasahara T, Kakiuchi C, Iwamoto K, Kawamura K, Ishihara H, Oka Y. Behavioral and gene expression analyses of Wfs1 knockout mice as a possible animal model of mood disorder. Neurosci Res. 2008; 61(2): 143–58.

Munshani S, Ibrahim EY, Domenicano I, Ehrlich BA. The Impact of Mutations in Wolframin on Psychiatric Disorders. Front Pediatr. 2021; 9: 718132.

Xavier J, Bourvis N, Tanet A, Ramos T, Perisse D, Marey I, Cohen D, Consoli A. Bipolar Disorder Type 1 in a 17-Year-Old Girl with Wolfram Syndrome. J Child Adolesc Psychopharmacol. 2016; 26(8): 750–755.

Szczepankiewicz D, Narożna B, Celichowski P, Sakrajda K, Kołodziejski P, Banach E, Zakowicz P, Pruszyńska-Oszmałek E, Pawlak J, Wiłkość M, Dmitrzak-Węglarz M, Skibińska M, Bejger A, Twarowska-Hauser J, Rybakowski JK, Nogowski L, Szczepankiewicz A. Genes involved in glucocorticoid receptor signaling affect susceptibility to mood disorders. World J Biol Psychiatry. 2021; 22(2): 149–160.

Kim J, Shim S, Choi SC, Han JK. A putative Xenopus Rho-GTPase activating protein (XrGAP) gene is expressed in the notochord and brain during early embryogenesis. Gene Expr Patterns. 2003; 3(2): 219–23.

Sekiguchi M, Sobue A, Kushima I, Wang C, Arioka Y, Kato H, Kodama A, Kubo H, Ito N, Sawahata M, Hada K, Ikeda R, Shinno M, Mizukoshi C, Tsujimura K, Yoshimi A, Ishizuka K, Takasaki Y, Kimura H, Xing J, Yu Y, Yamamoto M, Okada T, Shishido E, Inada T, Nakatochi M, Takano T, Kuroda K, Amano M, Aleksic B, Yamomoto T, Sakuma T, Aida T, Tanaka K, Hashimoto R, Arai M, Ikeda M, Iwata N, Shimamura T, Nagai T, Nabeshima T, Kaibuchi K, Yamada K, Mori D, Ozaki N. ARHGAP10, which encodes Rho GTPase-activating protein 10, is a novel gene for schizophrenia risk. Translational Psychiatry. 2020; 10(1): 247.

Hada K, Wulaer B, Nagai T, Itoh N, Sawahata M, Sobue A, Mizoguchi H, Mori D, Kushima I, Nabeshima T, Ozaki N, Yamada K. Mice carrying a schizophrenia-associated mutation of the Arhgap10 gene are vulnerable to the effects of methamphetamine treatment on cognitive function: association with morphological abnormalities in striatal neurons. Mol Brain. 2021; 14(1): 21.

Jope RS. A bimodal model of the mechanism of action of lithium. Mol Psychiatry. 1999; 4(1): 21–5.

Albert PR, Fiori LM. Transcriptional dys-regulation in anxiety and major depression: 5-HT1A gene promoter architecture as a therapeutic opportunity. Curr Pharm Des. 2014; 20(23): 3738–50.

Cox OH, Song HY, Garrison-Desany HM, Gadiwalla N, Carey JL, Menzies J, Lee RS. Characterization of glucocorticoid-induced loss of DNA methylation of the stress-response gene *Fkbp5* in neuronal cells. Epigenetics. 2021c; 16(12): 1377–1397.

Haussler MR, Whitfield GK, Haussler CA, Sabir MS, Khan Z, Sandoval R, Jurutka PW. 1,25-Dihydroxyvitamin D and Klotho: A Tale of Two Renal Hormones Coming of Age. Vitam Horm. 2016; 100: 165–230.

Khan WN, Teglund S, Bremer K, Hammarström S. The pregnancy-specific glycoprotein family of the immunoglobulin superfamily: identification of new members and estimation of family size. Genomics. 1992; 12(4): 780–7.

Luik RM, Lewis RS. New insights into the molecular mechanisms of store-operated Ca2+ signaling in T cells. Trends Mol Med. 2007r; 13(3): 103–7.

Miller AH. Depression and immunity: a role for T cells? Brain Behav Immun. 2010; 24(1): 1–8.

Vaeth M, Kahlfuss S, Feske S. CRAC Channels and Calcium Signaling in T Cell-Mediated Immunity. Trends Immunol. 2020; 41(10): 878–901.

Ramesh G, Jarzembowski L, Schwarz Y, Poth V, Konrad M, Knapp ML, Schwär G, Lauer AA, Grimm MOW, Alansary D, Bruns D, Niemeyer BA. A short isoform of STIM1 confers frequency-dependent synaptic enhancement. Cell Rep. 2021; 34(11): 108844.

Voros O, Panyi G, Hajdu P. Immune Synapse Residency of Orai1 Alters Ca^2+^ Response of T Cells. Int J Mol Sci. 2021; 22(21): 11514.

Hsiao CC, Sankowski R, Prinz M, Smolders J, Huitinga I, Hamann J. GPCRomics of Homeostatic and Disease-Associated Human Microglia. Front Immunol. 2021; 12: 674189.

Wise, Toby, J. Radua, E. Via, N. Cardoner, O. Abe, T. Adams, F. Amico, Y. Cheng, J. Cole, C. de Azevedo Marques Périco, et al. Common and distinct patterns of grey-matter volume alteration in major depression and bipolar disorder: evidence from a voxel-based meta-analysis. Molecular Psychiatry. 2017; 22(10): 1455–1463.

Johnston, Jennifer A. Y., Fei Wang, Jie Liu, Benjamin Blond, Amanda Wallace, Jiacheng Liu, Linda Spencer, Elizabeth Lippard, Kristin Purves, Angeli Landeros-Weisenberger, et al. Multimodal Neuroimaging of Frontolimbic Structure and Function Associated With Suicide Attempts in Adolescents and Young Adults With Bipolar Disorder. The American Journal of Psychiatry. 2017; 174(7): 667–675.

Schmaal, Lianne, Anne-Laura van Harmelen, Vasiliki Chatzi, Elizabeth Lippard, Yara Toenders, Lynnette Averill, Carolyn Mazure, and Hilary Blumberg. Imaging suicidal thoughts and behaviors: a comprehensive review of 2 decades of neuroimaging studies. Molecular Psychiatry. 2020; 25(2): 408–427.

Jabbi, Mbemba, Wade Weber, Jeffrey Welge, Fabiano Nery, Maxwell Tallman, Austin Gable, David Fleck, Elizabeth Lippard, Melissa Delbello, Caleb Adler, et al. Frontolimbic brain volume abnormalities in bipolar disorder with suicide attempts. Psychiatry Research. 2020b; 294:113516.

Salvadore, Giacomo, Brian Cornwell, Veronica Colon-Rosario, Richard Coppola, Christian Grillon, Carlos Zarate, and Husseini Manji. Increased anterior cingulate cortical activity in response to fearful faces: a neurophysiological biomarker that predicts rapid antidepressant response to ketamine. Biological Psychiatry. 2009; 65(4): 289–95.

McGrath, Callie. L., Mary Kelley, Paul Holtzheimer, Boadie Dunlop, W. Craighead, Alexandre Franco, R. Craddock, and Helen Mayberg. Toward a neuroimaging treatment selection biomarker for major depressive disorder. JAMA Psychiatry. 2013; 70(8): 821–829.

Riva-Posse, Patricio, K. Choi, P. Holtzheimer, A. Crowell, S. Garlow, J. Rajendra, C. McIntyre, R. Gross, and H. Mayberg. A connectomic approach for subcallosal cingulate deep brain stimulation surgery: prospective targeting in treatment-resistant depression. Molecular Psychiatry. 2018; 23(4): 843–849.

Galvalvy H, Zalsman G, Huang YY, Murphy L, Rosoklija G, Dwork AJ, Haghighi F, Arango V, Mann JJ. A pilot genome-wide association and gene expression array study of suicide with and without major depression. World J Biol Psychiatry. 2013; 14(8): 574–82.

Orozco-Solis R, Montellier E, Aguilar-Arnal L, Sato S, Vawter MP, Bunney BG, Bunney WE, Sassone-Corsi P. A Circadian Genomic Signature Common to Ketamine and Sleep Deprivation in the Anterior Cingulate Cortex. Biol Psychiatry. 2017; 82(5): 351–360.

Lutz PE, Tanti A, Gasecka A, Barnett-Burns S, Kim JJ, Zhou Y, Chen GG, Wakid M, Shaw M, Almeida D, Chay MA, Yang J, Larivière V, M’Boutchou MN, van Kempen LC, Yerko V, Prud’homme J, Davoli MA, Vaillancourt K, Théroux JF, Bramoullé A, Zhang TY, Meaney MJ, Ernst C, Côté D, Mechawar N, Turecki G Association of a History of Child Abuse With Impaired Myelination in the Anterior Cingulate Cortex: Convergent Epigenetic, Transcriptional, and Morphological Evidence. Am J Psychiatry. 2017; 174(12): 1185–1194.

Zhou Y, Lutz PE, Wang YC, Ragoussis J, Turecki G. Global long non-coding RNA expression in the rostral anterior cingulate cortex of depressed suicides. Transl Psychiatry. 2018; 8(1): 224.

Elhaik E, Zandi P. Dysregulation of the NF-κB pathway as a potential inducer of bipolar disorder. J Psychiatr Res. 2015; 70: 18–27.

Schiweck C, Valles-Colomer M, Arolt V, Müller N, Raes J, Wijkhuijs A, Claes S, Drexhage H, Vrieze E. Depression, and suicidality: A link to premature T helper cell aging and increased Th17 cells. Brain Behav Immun. 2020; 87: 603–609.

Traub N, Lichtstein D. The mood cycle hypothesis: possible involvement of steroid hormones in mood regulation by means of Na+, K+-ATPase inhibition. J Basic Clin Physiol Pharmacol. 2000;11(4): 375–94.

Blokland GAM et al. Sex-Dependent Shared and Nonshared Genetic Architecture Across Mood and Psychotic Disorders. Biol Psychiatry. 2022; 91(1): 102–117.

Glover V, O’Donnell KJ, O’Connor TG, Fisher J. Prenatal maternal stress, fetal programming, and mechanisms underlying later psychopathology-A global perspective. Dev Psychopathol. 2018; 30(3): 843–854.

Mechawar N., and J. Savitz. Neuropathology of mood disorders: do we see the stigmata of inflammation? Translational Psychiatry. 2016; 6(11): e946.

Giridharan, Vijayasree, Pavani Sayana, Omar Pinjari, Naveed Ahmad , Maria da Rosa, João Quevedo, and Tatiana Barichello. Postmortem evidence of brain inflammatory markers in bipolar disorder: a systematic review. Molecular Psychiatry. 2020; 25(1): 94–113.

Slavich G et al. Neural sensitivity to social rejection is associated with inflammatory responses to social stress. Proc Natl Acad Sci USA. 2010; 107(33): 14817–22.

Rivière JB, Mirzaa GM, O’Roak BJ, Beddaoui M, Alcantara D, Conway RL, St-Onge J, Schwartzentruber JA, Gripp KW, Nikkel SM, Worthylake T, Sullivan CT, Ward TR, Butler HE, Kramer NA, Albrecht B, Armour CM, Armstrong L, Caluseriu O, Cytrynbaum C, Drolet BA, Innes AM, Lauzon JL, Lin AE, Mancini GM, Meschino WS, Reggin JD, Saggar AK, Lerman-Sagie T, Uyanik G, Weksberg R, Zirn B, Beaulieu CL; Finding of Rare Disease Genes (FORGE) Canada Consortium; Majewski J, Bulman DE, O’Driscoll M, Shendure J, Graham JM Jr, Boycott KM, Dobyns WB. De novo germline and postzygotic mutations in AKT3, PIK3R2, and PIK3CA cause a spectrum of related megalencephaly syndromes. Nat Genet. 2012; 44(8):934–40

