## Supplementary material for "Brain transcriptomic signatures for mood disorders and suicide phenotypes: an anterior insula and subgenual ACC network postmortem study": Brain network transcriptomic signatures for mood disorders and suicide phenotypes_Supplementary Tables, Figures, and References2.docx

In the supplementary tables below, gray shaded results represent downregulated genes (negative Log2Foldchange values), whereas non-shaded results represent upregulated genes (positive Log2Foldchange values).

| **TABLE S1**  **TABLE S1A** | | |
| --- | --- | --- |
| Suicide Completion in Ant-Ins (Mood Disorders Only) | | |
| GeneName | log2FoldChange | q-adjusted |
| BAALC-AS1 | -1.39 | 0.0010 |
| FOSB | -2.11 | 0.0054 |
| SERPINA3 | -3.24 | 0.0054 |
| CHI3L1 | -1.82 | 0.0065 |
| AC145676.2 | -0.91 | 0.0084 |
| SLC39A14 | -0.89 | 0.0442 |
| **Table S1B** | | |
| Suicide completion in sgACC (Mood Disorders Only) | | |
| GeneName | log2FoldChange | q-adjusted |
| RP11-403A3.3 | -1.86 | 5.66E-07 |
| SERPINA3 | -3.24 | 5.66E-07 |
| CHI3L1 | -1.92 | 9.00E-07 |
| SFN | -2.14 | 3.62E-06 |
| SLC11A1 | -1.32 | 6.00E-06 |
| ICAM1 | -1.31 | 7.63E-06 |
| GBP2 | -1.26 | 1.27E-05 |
| TNFRSF1A | -0.82 | 1.37E-05 |
| BAG3 | -1.42 | 2.20E-05 |
| F3 | -1.04 | 2.20E-05 |
| IL1RL1 | -1.74 | 2.84E-05 |
| BAALC-AS1 | -0.75 | 6.25E-05 |
| HILPDA | -1.96 | 6.25E-05 |
| OSMR | -1.27 | 0.000101 |
| EMP1 | -1.38 | 0.000103 |
| KIAA0040 | -1.28 | 0.000106 |
| MTHFD2 | -0.81 | 0.000108 |
| PDPN | -1.07 | 0.000111 |
| SLC39A14 | -0.98 | 0.000111 |
| C1R | -0.78 | 0.000125 |
| SLC44A3 | -0.62 | 0.000125 |
| MYO1G | -1.23 | 0.000146 |
| MAP3K6 | -0.66 | 0.000156 |
| SIX4 | -0.96 | 0.000166 |
| SERPINA1 | -1.58 | 0.000175 |
| SERPINE1 | -1.10 | 0.000175 |
| MT1X | -1.50 | 0.000213 |
| RP11-473M20.16 | -1.26 | 0.000244 |
| HAMP | -1.25 | 0.000353 |
| SERPINH1 | -1.71 | 0.000353 |
| TNFRSF12A | -1.26 | 0.000405 |
| MYC | -0.94 | 0.000421 |
| FUT3 | -0.69 | 0.000431 |
| MGST1 | -0.45 | 0.000433 |
| TEAD3 | -0.76 | 0.000444 |
| CEBPD | -1.15 | 0.000501 |
| FPR1 | -1.14 | 0.000501 |
| GBP1 | -1.10 | 0.000501 |
| IL1R1 | -1.20 | 0.000501 |
| LINC01057 | -0.48 | 0.000564 |
| C1S | -0.59 | 0.000564 |
| CNN3 | -0.60 | 0.000564 |
| FGF2 | -0.68 | 0.000564 |
| HSPA7 | -1.70 | 0.000564 |
| DTNA | -0.55 | 0.000628 |
| BCL3 | -0.90 | 0.000658 |
| PLSCR1 | -0.87 | 0.000658 |
| SMAD1 | -0.25 | 0.000658 |
| DENND2D | -0.95 | 0.000747 |
| TPST1 | -0.49 | 0.000752 |
| CDKN1A | -1.43 | 0.000885 |
| ANGPTL4 | -1.36 | 0.00092 |
| CD163 | -0.67 | 0.000957 |
| LIMK2 | -0.64 | 0.001048 |
| YTHDF1 | -0.38 | 0.001234 |
| PNLDC1 | -1.07 | 0.001244 |
| ACTRT3 | -0.95 | 0.001422 |
| APOL6 | -0.38 | 0.001422 |
| IFITM2 | -1.17 | 0.001422 |
| LGALS3 | -0.53 | 0.001422 |
| TLR2 | -0.87 | 0.001422 |
| C1RL | -0.64 | 0.001448 |
| HMOX1 | -0.79 | 0.001448 |
| IL4R | -0.94 | 0.001448 |
| CD93 | -1.06 | 0.001473 |
| FCGR3A | -1.26 | 0.001528 |
| RP11-50D9.3 | -0.85 | 0.00186 |
| CEBPB | -0.51 | 0.001913 |
| CD14 | -1.08 | 0.001999 |
| PSTPIP2 | -0.65 | 0.0023 |
| A4GALT | -0.89 | 0.0023 |
| SOD2 | -0.35 | 0.002545 |
| PDLIM4 | -1.05 | 0.002983 |
| YBX3 | -0.82 | 0.002983 |
| GADD45A | -0.88 | 0.003079 |
| PIM1 | -0.65 | 0.003289 |
| ANO6 | -0.44 | 0.003376 |
| PLAC4 | -0.75 | 0.003639 |
| FCGR1A | -1.01 | 0.00367 |
| ADM | -1.78 | 0.003917 |
| MAOB | -0.38 | 0.003917 |
| ITPKC | -0.54 | 0.00415 |
| NAMPT | -0.55 | 0.004237 |
| BACE2 | -0.63 | 0.004328 |
| FERMT3 | -0.71 | 0.004588 |
| TIFA | -0.44 | 0.004588 |
| STOM | -0.46 | 0.004769 |
| C10orf10 | -1.10 | 0.004809 |
| GLIS3 | -0.62 | 0.004809 |
| MIR548AJ2 | -0.41 | 0.004809 |
| MT2A | -0.78 | 0.004809 |
| RP11-482G13.1 | -0.82 | 0.004809 |
| SBNO2 | -0.75 | 0.004819 |
| EMILIN2 | -0.65 | 0.004836 |
| OSMR-AS1 | -0.52 | 0.004836 |
| TUBB6 | -0.72 | 0.004997 |
| IFITM3 | -0.92 | 0.005059 |
| JAK3 | -0.55 | 0.005409 |
| SERPING1 | -0.58 | 0.005416 |
| ASAP3 | -0.39 | 0.005866 |
| PTPN2 | -0.19 | 0.005886 |
| RDH10 | -0.70 | 0.005886 |
| SORBS1 | -0.30 | 0.005886 |
| CP | -1.40 | 0.005896 |
| EMP3 | -0.60 | 0.006086 |
| ETV6 | -0.34 | 0.006086 |
| IFITM1 | -0.88 | 0.006086 |
| SOCS3 | -2.02 | 0.006274 |
| SLCO4A1 | -0.90 | 0.006389 |
| FAS | -0.56 | 0.006756 |
| HIF1A | -0.29 | 0.006981 |
| CLU | -0.35 | 0.007428 |
| CLEC2B | -0.54 | 0.007466 |
| RP11-211G3.2 | -0.87 | 0.007466 |
| STC1 | -1.04 | 0.007466 |
| CFI | -0.78 | 0.007491 |
| RP11-403A3.2 | -0.60 | 0.007537 |
| PLOD2 | -0.47 | 0.007671 |
| LSM6 | -0.26 | 0.007728 |
| SIGLEC9 | -0.59 | 0.007815 |
| AL773572.7 | -0.87 | 0.007954 |
| ALOXE3 | -1.00 | 0.008049 |
| CDK2 | -0.86 | 0.008049 |
| HSPB1 | -0.92 | 0.008049 |
| SLC16A3 | -0.74 | 0.008774 |
| GBP3 | -0.72 | 0.008781 |
| TMEM176B | -0.60 | 0.008936 |
| CTAGE6 | -1.09 | 0.009065 |
| ADAMTS9 | -0.95 | 0.00919 |
| CD59 | -0.26 | 0.009565 |
| PRRX1 | -0.39 | 0.009737 |
| AHCTF1 | -0.19 | 0.010316 |
| APOL4 | -0.30 | 0.01055 |
| G0S2 | -0.85 | 0.01055 |
| PLSCR4 | -0.43 | 0.01055 |
| TIPARP | -0.72 | 0.01055 |
| TRAF3IP2 | -0.31 | 0.01055 |
| STC2 | -1.19 | 0.010741 |
| CD44 | -1.32 | 0.010799 |
| TUBA1C | -0.48 | 0.010799 |
| MAN1C1 | -0.36 | 0.010991 |
| ZNF436-AS1 | -0.27 | 0.01116 |
| DSC2 | -0.39 | 0.0115 |
| LILRB3 | -0.41 | 0.011872 |
| RP11-106M7.1 | -0.42 | 0.011873 |
| FSTL1 | -0.54 | 0.011994 |
| MRVI1 | -0.52 | 0.012286 |
| BOC | -0.53 | 0.012413 |
| CASP1 | -0.72 | 0.012413 |
| PGAM2 | -1.25 | 0.012413 |
| STON1 | -0.88 | 0.012413 |
| TIMP1 | -0.63 | 0.012413 |
| ARID5A | -0.62 | 0.01257 |
| GNA14 | -0.56 | 0.01257 |
| NNMT | -0.95 | 0.01257 |
| RFX4 | -0.50 | 0.01257 |
| DSE | -0.32 | 0.012607 |
| GLRX | -0.45 | 0.012976 |
| IFI30 | -0.88 | 0.01324 |
| ADIRF-AS1 | -0.35 | 0.014055 |
| NFKB2 | -0.58 | 0.01449 |
| STEAP3 | -0.69 | 0.014582 |
| CISH | -1.18 | 0.014583 |
| MS4A6A | -0.39 | 0.014583 |
| RP11-274H2.5 | -0.75 | 0.014583 |
| SLCO4A1-AS1 | -0.78 | 0.015076 |
| TMBIM1 | -0.65 | 0.015981 |
| CRISPLD1 | -0.73 | 0.016063 |
| RP11-572O17.1 | -0.91 | 0.016346 |
| LINC01480 | -0.30 | 0.016437 |
| C2 | -0.37 | 0.016501 |
| PTPN22 | -0.36 | 0.016501 |
| RP13-20L14.6 | -0.46 | 0.016501 |
| RAB20 | -0.82 | 0.016549 |
| MR1 | -0.21 | 0.016807 |
| MRVI1-AS1 | -0.49 | 0.016807 |
| RAB13 | -0.47 | 0.017828 |
| STAT3 | -0.37 | 0.017828 |
| EZH2 | -0.28 | 0.01804 |
| AHCYL1 | -0.37 | 0.018132 |
| C1QTNF1 | -0.50 | 0.018132 |
| COL4A1 | -0.68 | 0.018132 |
| MFAP5 | -0.61 | 0.018132 |
| TRAF6 | -0.15 | 0.018132 |
| FCGR2A | -0.93 | 0.018167 |
| KCNE4 | -0.24 | 0.018167 |
| BMPR1A | -0.21 | 0.018179 |
| RDH10-AS1 | -0.92 | 0.01819 |
| THBD | -0.85 | 0.018724 |
| CASP7 | -0.46 | 0.01921 |
| SIGLEC14 | -0.76 | 0.019898 |
| PPP1R3B | -0.69 | 0.020232 |
| RP11-50D9.4 | -0.81 | 0.020365 |
| MT1M | -0.90 | 0.021105 |
| NOD1 | -0.43 | 0.021105 |
| RP11-304F15.3 | -0.43 | 0.021105 |
| SLC25A37 | -0.28 | 0.021105 |
| TNFRSF10D | -0.67 | 0.021105 |
| PHACTR2P1 | -0.56 | 0.021385 |
| SLC17A9 | -0.52 | 0.021577 |
| WWTR1 | -0.67 | 0.021613 |
| GPR4 | -0.58 | 0.021636 |
| MFHAS1 | -0.26 | 0.021636 |
| RP11-70J12.1 | -0.70 | 0.021636 |
| PPP1R18 | -0.48 | 0.021767 |
| LINC00869 | -0.18 | 0.022252 |
| NUPR1 | -0.46 | 0.022252 |
| SMTNL1 | -0.47 | 0.023367 |
| UNG | -0.21 | 0.023444 |
| ALPK1 | -0.23 | 0.023503 |
| TEAD2 | -0.63 | 0.023503 |
| ATXN7 | -0.14 | 0.024555 |
| AC063976.7 | -0.53 | 0.024808 |
| EPHA2 | -0.81 | 0.024808 |
| PPRC1 | -0.18 | 0.024808 |
| PYGL | -0.52 | 0.024808 |
| RHPN2 | -0.37 | 0.024808 |
| LTF | -1.53 | 0.024876 |
| TNFSF13B | -0.40 | 0.024876 |
| FAM20C | -0.46 | 0.024904 |
| THBS1 | -0.67 | 0.024904 |
| LINC01532 | -0.64 | 0.025127 |
| PDLIM1 | -0.83 | 0.02547 |
| IER5L | -0.52 | 0.025545 |
| AEBP1 | -0.87 | 0.025662 |
| AQP4 | -0.52 | 0.025662 |
| ASPH | -0.20 | 0.025662 |
| DTX3L | -0.42 | 0.025662 |
| NSMAF | -0.13 | 0.025662 |
| PNRC1 | -0.23 | 0.025662 |
| RGS16 | -0.81 | 0.025662 |
| RP11-244H3.1 | -0.35 | 0.025662 |
| SQRDL | -0.48 | 0.025662 |
| TGM2 | -0.70 | 0.025662 |
| TAP1 | -0.52 | 0.025688 |
| RHOC | -0.37 | 0.026709 |
| CSTB | -0.19 | 0.026831 |
| IL1B | -1.26 | 0.026831 |
| DDIT4L | -0.58 | 0.027646 |
| RIN3 | -0.37 | 0.028413 |
| ZFP36 | -1.33 | 0.028823 |
| MYZAP | -0.76 | 0.029168 |
| ARHGEF35 | -0.68 | 0.029223 |
| SELP | -0.63 | 0.029398 |
| C4A-AS1 | -0.56 | 0.029625 |
| C4B-AS1 | -0.56 | 0.029625 |
| SPHK1 | -0.64 | 0.029645 |
| RP11-274H2.2 | -0.32 | 0.029979 |
| ATP6V1B1 | -0.71 | 0.030165 |
| S100A10 | -0.74 | 0.030165 |
| HLA-F-AS1 | -0.33 | 0.030194 |
| IL15RA | -0.64 | 0.030194 |
| IL1RAP | -0.23 | 0.030345 |
| DNAJB1 | -0.87 | 0.030395 |
| RP11-122G18.12 | -0.69 | 0.0306 |
| BHLHE40 | -0.38 | 0.030667 |
| P4HA2 | -0.37 | 0.030667 |
| TMEM132E | -0.49 | 0.030667 |
| TYMP | -0.58 | 0.030667 |
| FEM1C | -0.27 | 0.031134 |
| CCND2-AS1 | -0.63 | 0.031309 |
| POM121L9P | -0.67 | 0.031559 |
| RELL1 | -0.49 | 0.031743 |
| S100A11 | -0.73 | 0.031935 |
| ANXA2 | -0.65 | 0.031942 |
| LINC00963 | -0.24 | 0.031942 |
| MPZL2 | -0.65 | 0.032269 |
| STEAP4 | -0.28 | 0.032458 |
| SEMA4B | -0.26 | 0.03252 |
| VEGFA | -0.85 | 0.033613 |
| RP1-111D6.3 | -0.92 | 0.033864 |
| IFNLR1 | -0.58 | 0.034401 |
| TM4SF1 | -0.81 | 0.034401 |
| CD247 | -0.38 | 0.035039 |
| NAV2 | -0.27 | 0.035039 |
| CTC-444N24.8 | -0.28 | 0.035165 |
| TMEM176A | -0.44 | 0.035544 |
| VASN | -0.51 | 0.035544 |
| RBM27 | -0.12 | 0.035938 |
| ACSS3 | -0.26 | 0.03604 |
| ZNF292 | -0.10 | 0.036071 |
| RP11-798K3.2 | -0.44 | 0.036773 |
| GJA4 | -0.51 | 0.037179 |
| EPHA1-AS1 | -0.45 | 0.037407 |
| DDIT4 | -0.72 | 0.038126 |
| TCAF2 | -0.31 | 0.038126 |
| PDZK1 | -0.34 | 0.03836 |
| RUNX1 | -0.41 | 0.03836 |
| PML | -0.31 | 0.038397 |
| SEC24A | -0.11 | 0.039171 |
| RAC2 | -0.61 | 0.039633 |
| CHSY1 | -0.26 | 0.042002 |
| NEAT1 | -0.84 | 0.042297 |
| TGFB2 | -0.37 | 0.042408 |
| SLC10A1 | -0.62 | 0.04318 |
| RP11-274H2.3 | -0.42 | 0.043428 |
| FABP5 | -0.35 | 0.043496 |
| SYNM | -0.33 | 0.043496 |
| CASP4 | -0.41 | 0.043729 |
| S1PR3 | -0.31 | 0.043729 |
| PIRT | -0.95 | 0.0441 |
| RP11-61L23.2 | -0.66 | 0.044521 |
| BMPR1B | -0.37 | 0.044952 |
| GFPT2 | -0.28 | 0.044952 |
| GADD45G | -0.77 | 0.045023 |
| EDN1 | -0.69 | 0.047743 |
| MDK | -0.41 | 0.047762 |
| MAP3K14-AS1 | -0.23 | 0.048166 |
| TRIP10 | -0.69 | 0.048203 |
| HSD3B7 | -0.27 | 0.048524 |
| MAFF | -0.61 | 0.048525 |
| ANP32E | -0.20 | 0.048776 |
| FAM157A | -0.64 | 0.048776 |
| LINC00996 | 0.57 | 0.00092 |
| NAA40 | 0.15 | 0.005409 |
| KLHL3 | 0.16 | 0.006086 |
| NPAS4 | 1.96 | 0.007728 |
| COL22A1 | 0.52 | 0.010622 |
| RERGL | 0.81 | 0.01223 |
| NUDT16L1 | 0.21 | 0.012413 |
| ST8SIA2 | 0.76 | 0.013765 |
| RNU4-1 | 0.52 | 0.018179 |
| EDN3 | 0.65 | 0.01921 |
| AC005592.2 | 0.32 | 0.023367 |
| BDNF | 0.56 | 0.030194 |
| SMUG1 | 0.2 | 0.030395 |
| ABCG2 | 0.61 | 0.032191 |
| CRHBP | 0.51 | 0.032558 |
| KIRREL3-AS2 | 0.36 | 0.039855 |
| FAM13B | 0.16 | 0.043496 |
| SPRY4 | 0.33 | 0.044314 |
| CRH | 0.79 | 0.044532 |

|  | **TABLE S2** | | |
| --- | --- | --- | --- |
|  | **Table S2A** | | |
|  | Longevity in Ant-Ins (Mood Disorders & Controls) | | |
|  | GeneName | log2FoldChange | q-adjusted |
|  | MTND1P23 | -2.89 | 1.20E-06 |
|  | ARPC5 | -0.19 | 0.000526 |
|  | GSG1 | -0.39 | 0.000723 |
|  | TCEB1 | -0.18 | 0.008443 |
|  | RP11-1263C18.1 | -0.23 | 0.009502 |
|  | HSBP1 | -0.19 | 0.013489 |
|  | NDUFS4 | -0.22 | 0.016313 |
|  | SET | -0.14 | 0.016313 |
|  | METTL9 | -0.12 | 0.017446 |
|  | LINC01102 | -0.26 | 0.017512 |
|  | LMO4 | -0.24 | 0.018691 |
|  | MSANTD3-TMEFF1 | -0.24 | 0.021389 |
|  | CXADR | -0.23 | 0.022835 |
|  | CRK | -0.11 | 0.023006 |
|  | ARPC2 | -0.17 | 0.028745 |
|  | CHCHD3 | -0.13 | 0.028745 |
|  | PDCD2 | -0.12 | 0.028745 |
|  | RP11-271F18.4 | -0.28 | 0.028745 |
|  | SMIM10L1 | -0.19 | 0.028745 |
|  | PRKRA | -0.12 | 0.028935 |
|  | RALA | -0.13 | 0.028935 |
|  | DLGAP1-AS4 | -0.36 | 0.032106 |
|  | B3GNT2 | -0.26 | 0.032687 |
|  | DR1 | -0.17 | 0.032687 |
|  | FAM19A1 | -0.32 | 0.032687 |
|  | GVINP1 | -0.41 | 0.032687 |
|  | ST8SIA2 | -0.53 | 0.032687 |
|  | XRCC3 | 0.25 | 0.032687 |
|  | NDUFB6 | -0.19 | 0.033656 |
|  | USMG5 | -0.26 | 0.033656 |
|  | RP11-563K23.1 | -0.29 | 0.034034 |
|  | AARD | -0.39 | 0.03665 |
|  | BEX5 | -0.23 | 0.03665 |
|  | BZW1 | -0.13 | 0.03665 |
|  | C19orf81 | -0.40 | 0.03665 |
|  | CDC42 | -0.17 | 0.03665 |
|  | FRMPD2 | -0.43 | 0.03665 |
|  | NRBF2 | -0.18 | 0.03665 |
|  | SDCBP | -0.11 | 0.03665 |
|  | SMIM8 | -0.15 | 0.03665 |
|  | TTR | -1.38 | 0.03665 |
|  | UBE2F | -0.16 | 0.03665 |
|  | VPS29 | -0.14 | 0.03665 |
|  | BMP2 | -0.32 | 0.038298 |
|  | FRG1 | -0.13 | 0.038298 |
|  | RPS29 | -0.27 | 0.038298 |
|  | SDHAF3 | -0.18 | 0.038298 |
|  | UBXN2B | -0.17 | 0.038298 |
|  | PSMA2 | -0.14 | 0.039923 |
|  | CASC15 | -0.19 | 0.03998 |
|  | ADRA2A | -0.35 | 0.040463 |
|  | MYL12B | -0.18 | 0.040463 |
|  | STMN2 | -0.31 | 0.040463 |
|  | BLMH | -0.11 | 0.041902 |
|  | LSM3 | -0.19 | 0.042531 |
|  | PRPS2 | -0.27 | 0.042531 |
|  | ARNTL | -0.18 | 0.043187 |
|  | C8orf34 | -0.20 | 0.043187 |
|  | CCDC117 | -0.15 | 0.043187 |
|  | COX7C | -0.22 | 0.043187 |
|  | PCDHB2 | -0.29 | 0.043187 |
|  | PPEF1 | -0.32 | 0.043187 |
|  | SELT | -0.10 | 0.043187 |
|  | SIAH2 | -0.18 | 0.043187 |
|  | SLN | -0.61 | 0.043187 |
|  | EIF2S2 | -0.14 | 0.043683 |
|  | PTENP1 | -0.22 | 0.044001 |
|  | SERBP1 | -0.10 | 0.044001 |
|  | UBE2B | -0.14 | 0.044643 |
|  | DYRK2 | -0.25 | 0.046158 |
|  | C7orf25 | -0.30 | 0.046343 |
|  | MARCH1 | -0.23 | 0.046363 |
|  | ATP5E | -0.23 | 0.046516 |
|  | ATP5J | -0.20 | 0.046516 |
|  | CCL2 | -1.35 | 0.046516 |
|  | BZW2 | -0.20 | 0.048205 |
|  | CCDC167 | -0.22 | 0.048773 |
|  | KLRK1 | -0.51 | 0.048773 |
|  | ATP5I | -0.32 | 0.049094 |
|  | GMEB1 | -0.10 | 0.049094 |
|  | MMADHC | -0.14 | 0.049094 |
|  | TRBC2 | -0.54 | 0.049094 |
|  | FUK | 0.22 | 0.001333 |
|  | UAP1L1 | 0.30 | 0.002421 |
|  | UAP1L1 | 0.30 | 0.002421 |
|  | WFS1 | 0.37 | 0.008443 |
|  | WFS1 | 0.37 | 0.008443 |
|  | LTBP3 | 0.28 | 0.009502 |
|  | NUMA1 | 0.16 | 0.013489 |
|  | IQCA1 | 0.32 | 0.014883 |
|  | SPPL2B | 0.19 | 0.016313 |
|  | NAGLU | 0.25 | 0.017446 |
|  | AGFG2 | 0.21 | 0.020314 |
|  | TAF6L | 0.19 | 0.023006 |
|  | RHCG | 0.28 | 0.024415 |
|  | CFAP70 | 0.34 | 0.028745 |
|  | TMPRSS5 | 0.46 | 0.028745 |
|  | PAQR7 | 0.21 | 0.028935 |
|  | PIDD1 | 0.20 | 0.034034 |
|  | CPSF7 | 0.08 | 0.03665 |
|  | DNASE2 | 0.24 | 0.03665 |
|  | GREB1 | 0.19 | 0.03665 |
|  | IGDCC4 | 0.44 | 0.03665 |
|  | OPLAH | 0.48 | 0.03665 |
|  | PC | 0.22 | 0.03665 |
|  | PCDHGB1 | 0.44 | 0.03665 |
|  | PCSK4 | 0.30 | 0.03665 |
|  | PLA2G6 | 0.17 | 0.03665 |
|  | SGSH | 0.23 | 0.03665 |
|  | SSH3 | 0.22 | 0.03665 |
|  | WDR81 | 0.19 | 0.03665 |
|  | BRAT1 | 0.19 | 0.038298 |
|  | MEGF6 | 0.41 | 0.038298 |
|  | PCDHGA2 | 0.51 | 0.038298 |
|  | PLXNB2 | 0.23 | 0.038298 |
|  | RECQL5 | 0.11 | 0.039439 |
|  | ACCS | 0.30 | 0.040271 |
|  | VWCE | 0.23 | 0.040271 |
|  | SLC34A3 | 0.40 | 0.040395 |
|  | TMEM79 | 0.25 | 0.040463 |
|  | ITGB4 | 0.72 | 0.041902 |
|  | XYLT2 | 0.16 | 0.041902 |
|  | FAM90A1 | 0.26 | 0.041957 |
|  | DPP7 | 0.20 | 0.042531 |
|  | PQLC2 | 0.17 | 0.042531 |
|  | PYGB | 0.17 | 0.042531 |
|  | UNG | 0.28 | 0.042531 |
|  | IVD | 0.12 | 0.043187 |
|  | KCNN3 | 0.40 | 0.043187 |
|  | PLEC | 0.19 | 0.043187 |
|  | PARP3 | 0.15 | 0.043398 |
|  | ING5 | 0.10 | 0.043482 |
|  | GYS1 | 0.21 | 0.044001 |
|  | LINC00499 | 0.49 | 0.044001 |
|  | GMPR | 0.52 | 0.046363 |
|  | TFAP4 | 0.15 | 0.046363 |
|  | TPD52L2 | 0.09 | 0.046363 |
|  | CRB2 | 0.45 | 0.046516 |
|  | UNC93B1 | 0.36 | 0.046516 |
|  | CNNM3 | 0.14 | 0.048773 |
|  | CRYL1 | 0.25 | 0.048773 |
|  | KLHL21 | 0.18 | 0.048773 |
|  | KDM3A | 0.15 | 0.048944 |
|  | LFNG | 0.54 | 0.049094 |
|  | TPCN1 | 0.30 | 0.049094 |
|  | SLC4A11 | 0.48 | 0.049783 |
|  | Longevity in Ant-Ins (Mood Disorders only) | | |
|  | GeneName | log2FoldChange | q-adjusted |
|  | IQCA1 | -0.43 | 0.0007 |
|  | C4B | -0.98 | 0.0157 |
|  | GMPR | -0.78 | 0.0157 |
|  | WFS1 | -0.44 | 0.0157 |
|  | IGDCC4 | -0.56 | 0.0162 |
|  | TMPRSS5 | -0.58 | 0.0162 |
|  | SLC46A1 | -0.31 | 0.0172 |
|  | IGSF11 | -0.31 | 0.0174 |
|  | CCDC40 | -0.31 | 0.0176 |
|  | KDM3A | -0.23 | 0.0176 |
|  | HPR | -0.87 | 0.0192 |
|  | RHCG | -0.36 | 0.0192 |
|  | TPP1 | -0.48 | 0.0192 |
|  | BBS2 | -0.32 | 0.0202 |
|  | CTD-2353F22.1 | -1.25 | 0.0202 |
|  | LHFPL1 | -0.93 | 0.0202 |
|  | LINC00092 | -0.54 | 0.0202 |
|  | UNG | -0.39 | 0.0202 |
|  | CLU | -0.47 | 0.0204 |
|  | KCNN3 | -0.55 | 0.0204 |
|  | CFAP70 | -0.42 | 0.0209 |
|  | ACCS | -0.39 | 0.0212 |
|  | FUK | -0.22 | 0.0212 |
|  | GPR143 | -0.69 | 0.0212 |
|  | AGFG2 | -0.26 | 0.0214 |
|  | CTSH | -0.63 | 0.0234 |
|  | DNAH7 | -0.37 | 0.0234 |
|  | OPLAH | -0.62 | 0.0234 |
|  | LPIN1 | -0.21 | 0.0242 |
|  | DOCK7 | -0.32 | 0.0247 |
|  | LPP-AS2 | -0.48 | 0.0247 |
|  | H6PD | -0.31 | 0.0262 |
|  | TSPAN33 | -0.22 | 0.0262 |
|  | PLK5 | -0.42 | 0.0268 |
|  | RP5-858L17.1 | -0.34 | 0.0268 |
|  | MCCC2 | -0.24 | 0.0277 |
|  | DHX58 | -0.35 | 0.0288 |
|  | AC005336.4 | -0.96 | 0.0291 |
|  | PLIN4 | -0.97 | 0.0291 |
|  | C4A | -0.88 | 0.0291 |
|  | CD109 | -0.46 | 0.0291 |
|  | CTC-498M16.2 | -0.50 | 0.0291 |
|  | GYS1 | -0.28 | 0.0291 |
|  | PAMR1 | -0.62 | 0.0291 |
|  | PIK3IP1 | -0.33 | 0.0291 |
|  | ZNRF3 | -0.64 | 0.0291 |
|  | MIR3622A | -0.54 | 0.0303 |
|  | OSBPL11 | -0.43 | 0.0303 |
|  | PCDHGA2 | -0.63 | 0.0311 |
|  | LINC00499 | -0.62 | 0.0311 |
|  | DNASE2 | -0.30 | 0.0314 |
|  | IVD | -0.15 | 0.0314 |
|  | PHYHD1 | -0.61 | 0.0314 |
|  | RP11-159D12.2 | -0.27 | 0.0314 |
|  | RP11-437J2.3 | -0.38 | 0.0314 |
|  | SPARCL1 | -0.35 | 0.0314 |
|  | CPSF7 | -0.10 | 0.0329 |
|  | UAP1L1 | -0.29 | 0.0329 |
|  | NTRK2 | -0.46 | 0.0345 |
|  | AC069368.3 | -0.75 | 0.0358 |
|  | BMPR1B | -0.70 | 0.0361 |
|  | HVCN1 | -0.47 | 0.0361 |
|  | ACOX1 | -0.25 | 0.0363 |
|  | UNC93B1 | -0.43 | 0.0363 |
|  | ACACB | -0.57 | 0.0364 |
|  | ITGB4 | -0.89 | 0.0364 |
|  | SLC22A5 | -0.30 | 0.0364 |
|  | BMP2K | -0.40 | 0.0366 |
|  | SEMA4B | -0.41 | 0.0377 |
|  | NUMA1 | -0.18 | 0.0381 |
|  | LGI4 | -0.32 | 0.0384 |
|  | RP11-806O11.1 | -0.61 | 0.0387 |
|  | NOTCH2NL | -0.49 | 0.0392 |
|  | AQP4 | -0.73 | 0.0401 |
|  | CTD-2353F22.2 | -1.09 | 0.0401 |
|  | DHODH | -0.18 | 0.0401 |
|  | KREMEN1 | -0.29 | 0.0401 |
|  | LTBP3 | -0.28 | 0.0401 |
|  | PCDHGA3 | -0.72 | 0.0401 |
|  | SORCS2 | -0.42 | 0.0401 |
|  | SPPL2B | -0.21 | 0.0401 |
|  | ACOT11 | -0.37 | 0.0421 |
|  | ARHGAP5-AS1 | -0.37 | 0.0421 |
|  | TCF7 | -0.42 | 0.0422 |
|  | ADAMTSL5 | -0.46 | 0.0423 |
|  | NPFFR1 | -0.47 | 0.0423 |
|  | PHKG1 | -0.52 | 0.0423 |
|  | SLC14A1 | -1.03 | 0.0423 |
|  | TMCO4 | -0.34 | 0.0424 |
|  | BCL2 | -0.33 | 0.0436 |
|  | CRYL1 | -0.32 | 0.0436 |
|  | MAOB | -0.21 | 0.0436 |
|  | RHPN2 | -0.58 | 0.0436 |
|  | RP6-201G10.2 | -0.47 | 0.0446 |
|  | RP11-517I3.1 | -0.66 | 0.0447 |
|  | SLC4A11 | -0.53 | 0.0447 |
|  | ABCC11 | -0.50 | 0.0461 |
|  | PLXNB2 | -0.28 | 0.0468 |
|  | C10orf105 | -1.00 | 0.0468 |
|  | FUT10 | -0.43 | 0.0468 |
|  | GPT2 | -0.39 | 0.0468 |
|  | LCTL | -0.53 | 0.0468 |
|  | SLC34A3 | -0.49 | 0.0468 |
|  | SLC18B1 | -0.40 | 0.0479 |
|  | HHATL | -0.52 | 0.0479 |
|  | PRDM16 | -0.58 | 0.0483 |
|  | ATP13A4 | -0.67 | 0.0484 |
|  | HSPBAP1 | -0.21 | 0.0484 |
|  | MUC1 | -0.57 | 0.0484 |
|  | AC004019.13 | -0.58 | 0.0488 |
|  | AK4 | -0.26 | 0.0488 |
|  | BCAR3 | -0.38 | 0.0488 |
|  | EEF2K | -0.29 | 0.0488 |
|  | FOSB | -1.35 | 0.0488 |
|  | MROH7 | -0.39 | 0.0488 |
|  | NEBL | -0.28 | 0.0488 |
|  | PLCG1-AS1 | -0.54 | 0.0488 |
|  | PLEKHG4 | -0.36 | 0.0488 |
|  | RP11-106M7.1 | -0.40 | 0.0488 |
|  | RP11-1072A3.3 | -0.34 | 0.0488 |
|  | ZNF491 | -0.32 | 0.0488 |
|  | RP11-388C12.8 | -0.41 | 0.0489 |
|  | ADHFE1 | -0.44 | 0.0489 |
|  | ALS2CR12 | -0.28 | 0.0489 |
|  | GREB1 | -0.21 | 0.0489 |
|  | TPD52L1 | -0.53 | 0.0489 |
|  | VMAC | -0.24 | 0.0489 |
|  | CYP4F11 | -0.67 | 0.0494 |
|  | EPHX1 | -0.56 | 0.0496 |
|  | MTND1P23 | 3.26 | 1.20E-09 |
|  | MTCO1P12 | 2.38 | 3.87E-05 |
|  | ARPC5 | 0.22 | 0.0157 |
|  | SMIM10L1 | 0.26 | 0.0157 |
|  | ST8SIA2 | 0.76 | 0.0157 |
|  | RALA | 0.16 | 0.0176 |
|  | CASC15 | 0.27 | 0.0192 |
|  | ARPC2 | 0.22 | 0.0202 |
|  | BZW1 | 0.16 | 0.0202 |
|  | CSMD2 | 0.24 | 0.0202 |
|  | METTL9 | 0.16 | 0.0202 |
|  | SET | 0.17 | 0.0202 |
|  | NKAIN2 | 0.26 | 0.0210 |
|  | DR1 | 0.22 | 0.0234 |
|  | CELSR1 | 0.54 | 0.0242 |
|  | HSBP1 | 0.22 | 0.0259 |
|  | TMEM64 | 0.22 | 0.0268 |
|  | CHCHD3 | 0.15 | 0.0276 |
|  | RP11-271F18.4 | 0.33 | 0.0288 |
|  | SERBP1 | 0.14 | 0.0288 |
|  | LMO4 | 0.27 | 0.0291 |
|  | RP11-136K7.2 | 0.61 | 0.0291 |
|  | SOBP | 0.25 | 0.0291 |
|  | ANO4 | 0.26 | 0.0303 |
|  | CDH6 | 0.30 | 0.0310 |
|  | SDCBP | 0.14 | 0.0310 |
|  | H19 | 1.24 | 0.0329 |
|  | C8orf34 | 0.34 | 0.0330 |
|  | LGALSL | 0.21 | 0.0364 |
|  | PDCD2 | 0.15 | 0.0364 |
|  | USP12 | 0.27 | 0.0378 |
|  | PKP4 | 0.24 | 0.0384 |
|  | NDUFS4 | 0.23 | 0.0401 |
|  | SEMA3C | 0.37 | 0.0401 |
|  | UBXN2B | 0.21 | 0.0401 |
|  | NPPA | 0.60 | 0.0421 |
|  | RNF152 | 0.33 | 0.0421 |
|  | STMN2 | 0.38 | 0.0421 |
|  | ENPP6 | 0.47 | 0.0423 |
|  | SLC35G1 | 0.29 | 0.0423 |
|  | PTHLH | 0.36 | 0.0424 |
|  | SMARCE1 | 0.10 | 0.0436 |
|  | TC2N | 0.35 | 0.0436 |
|  | MEST | 0.24 | 0.0447 |
|  | SLN | 0.71 | 0.0447 |
|  | GNB4 | 0.25 | 0.0468 |
|  | GSG1 | 0.33 | 0.0468 |
|  | LINC00662 | 0.18 | 0.0468 |
|  | MARCH1 | 0.27 | 0.0468 |
|  | TCEB1 | 0.17 | 0.0479 |
|  | RAPGEF4 | 0.32 | 0.0483 |
|  | BLMH | 0.14 | 0.0484 |
|  | MAML3 | 0.24 | 0.0484 |
|  | MSANTD3-TMEFF1 | 0.24 | 0.0488 |
|  | NFU1 | 0.19 | 0.0488 |
|  | RP11-1263C18.1 | 0.24 | 0.0488 |
|  | YWHAB | 0.22 | 0.0488 |
|  | CXADR | 0.24 | 0.0489 |
|  | CDC42 | 0.19 | 0.0489 |
|  | MTPN | 0.21 | 0.0489 |
|  | EGFEM1P | 0.40 | 0.0494 |
|  | PPEF1 | 0.38 | 0.0494 |
|  | TYRP1 | 0.49 | 0.0496 |
|  | ZC3H15 | 0.21 | 0.0500 |
|  | **Table S2B** |  |  |
| Longevity in subg-ACC (Mood Disorders & Controls) | | | |
|  | GeneName | log2FoldChange | q-adjusted |
|  | ARHGAP10 | -0.33 | 0.0215 |
|  | CPXM1 | -0.58 | 0.0362 |
|  | PTGER4 | -0.45 | 0.0362 |
|  | RANBP17 | -0.21 | 0.0362 |
|  | ADGRG6 | -0.38 | 0.0426 |
|  | TNFRSF10A | -0.45 | 0.0426 |
|  | ISG15 | -0.36 | 0.0435 |
|  | MALL | -0.34 | 0.0435 |
|  | RNF152 | -0.25 | 0.0435 |
|  | SOX4 | -0.22 | 0.0435 |
|  | TH | -0.83 | 0.0435 |
|  | ZFP37 | -0.15 | 0.0435 |
|  | ACCS | 0.20 | 0.0435 |
|  | ROCK1P1 | 0.76 | 0.0435 |
|  | Longevity in subg-ACC (Mood Disorders only) | | |
|  | GeneName | log2FoldChange | q-adjusted |
|  | ARHGAP10 | -0.45 | 0.0037 |
|  | AC012146.7 | -0.35 | 0.0164 |
|  | FBXO18 | -0.11 | 0.0240 |
|  | RACGAP1 | -0.18 | 0.0240 |
|  | USP12 | -0.17 | 0.0240 |
|  | COL11A1 | -0.22 | 0.0400 |
|  | FAM184B | -0.20 | 0.0400 |
|  | WWC2 | -0.17 | 0.0400 |
|  | ZNF23 | -0.13 | 0.0400 |
|  | B4GALT2 | -0.26 | 0.0485 |
|  | ROCK1P1 | 1.22 | 0.0014 |
|  | RP11-474N24.6 | 0.32 | 0.0164 |
|  | CTC-591M7.1 | 0.25 | 0.0240 |
|  | PRELP | 0.37 | 0.0240 |
|  | RP11-138A9.1 | 0.32 | 0.0240 |
|  | RP11-138A9.2 | 0.26 | 0.0240 |
|  | RP11-269G24.7 | 0.47 | 0.0386 |
|  | C12orf60 | 0.17 | 0.0400 |
|  | CTD-2014D20.1 | 0.35 | 0.0400 |
|  | RP11-265O12.1 | 0.27 | 0.0400 |
|  | RP11-214K3.21 | 0.26 | 0.0485 |
|  | RP11-283I3.6 | 0.18 | 0.0485 |

| **Table S3**  **Table S3A** | | |
| --- | --- | --- |
| Longevity in Ant-Ins (Controls Only) | | |
| GeneName | log2FoldChange | q-adjusted |
| RP11-638I2.6 | -1.24 | 8.03E-06 |
| RP4-738P15.6 | -1.39 | 0.0019 |
| CTB-43P18.1 | -0.64 | 0.0030 |
| HCG25 | -0.82 | 0.0030 |
| RP1-257A7.5 | -0.69 | 0.0030 |
| RP11-215P8.2 | -0.60 | 0.0031 |
| PSMA2 | -0.23 | 0.0032 |
| RPL9 | -0.89 | 0.0044 |
| PET100 | -0.47 | 0.0081 |
| COL6A3 | -1.05 | 0.0084 |
| MT-TS1 | -1.10 | 0.0084 |
| RP11-603J24.17 | -0.93 | 0.0102 |
| H2BFM | -0.59 | 0.0137 |
| C19orf81 | -0.58 | 0.0148 |
| RPS29 | -0.37 | 0.0256 |
| AC005944.2 | -0.71 | 0.0262 |
| SLIRP | -0.35 | 0.0278 |
| AC009133.12 | -0.53 | 0.0323 |
| ATP5I | -0.49 | 0.0349 |
| AC007192.6 | -1.45 | 0.0388 |
| CTC-490E21.14 | -0.59 | 0.0395 |
| C10orf67 | -0.36 | 0.0401 |
| CTD-2562J17.7 | -1.06 | 0.0454 |
| RP11-361L15.3 | -0.41 | 0.0454 |
| GSG1 | -0.48 | 0.0486 |
| FAM60DP | 3.26 | 3.64E-06 |
| RP11-325O24.5 | 3.82 | 3.64E-06 |
| RPSAP48 | 3.14 | 1.33E-05 |
| PSG2 | 3.02 | 3.81E-05 |
| FGF23 | 2.44 | 5.31E-05 |
| LPAR4 | 2.56 | 0.0002 |
| S1PR2 | 1.50 | 0.0003 |
| RP11-44M6.3 | 2.60 | 0.0003 |
| MIR3648-1 | 0.63 | 0.0005 |
| MIR3648-2 | 0.63 | 0.0005 |
| NPM1P40 | 1.99 | 0.0005 |
| RP11-715J22.2 | 1.78 | 0.0005 |
| SHOX | 1.60 | 0.0005 |
| RP11-38O23.4 | 2.03 | 0.0006 |
| AC009404.2 | 1.91 | 0.0015 |
| FOXP4 | 0.31 | 0.0019 |
| WT1-AS | 2.16 | 0.0019 |
| RP11-87N24.3 | 1.93 | 0.0026 |
| MIR663AHG | 0.58 | 0.0027 |
| LRRC69 | 1.54 | 0.0030 |
| LINC00273 | 0.54 | 0.0031 |
| BMP8B | 0.59 | 0.0048 |
| RP1-17K7.2 | 1.31 | 0.0075 |
| ZNF302 | 0.68 | 0.0137 |
| MIR6087 | 0.86 | 0.0152 |
| RP11-217O12.1 | 1.53 | 0.0177 |
| CTD-2527I21.4 | 0.64 | 0.0211 |
| ZNF121 | 0.38 | 0.0227 |
| MIR3687-1 | 0.43 | 0.0256 |
| MIR3687-2 | 0.43 | 0.0256 |
| CTC-204F22.1 | 0.70 | 0.0298 |
| AGAP7P | 1.39 | 0.0405 |
| SYNDIG1L | 1.74 | 0.0447 |
| NTN3 | 0.49 | 0.0494 |
| **Table S3B** |  |  |
| Longevity in Subg-ACC (Controls Only) | | |

| Gene Name | Log2 Fold Change | q-adjusted |
| --- | --- | --- |
| SELE | -2.086484395 | 0.00272583 |
| TNFRSF10A | -0.755720757 | 0.00272583 |
| TMEM45B | -0.356020359 | 0.0053495 |
| SEMA3F | -0.59504703 | 0.00898067 |
| ADGRL4 | -0.45364579 | 0.01710602 |
| PUDP | -0.342556881 | 0.01971902 |
| VASP | -0.369964092 | 0.03062816 |
| PTGER4 | -0.695458526 | 0.03530824 |
| HSPA1A | -1.485909733 | 0.0381877 |
| HSPB1 | -1.179515039 | 0.0381877 |
| PLA1A | -1.042718997 | 0.0381877 |
| PLEKHG1 | -0.288379801 | 0.0381877 |
| SOCS3 | -1.618596536 | 0.0464215 |
| ADAMTS1 | -0.959752544 | 0.04833743 |
| DNAJB1 | -1.126933559 | 0.04876992 |
| ICAM2 | -0.393988889 | 0.04876992 |
| NOS3 | -0.495096445 | 0.04876992 |
| ORAI1 | -0.338937288 | 0.04876992 |
| AF131216.6 | -0.205331634 | 0.04887092 |
| CNN2 | -0.45579366 | 0.04887092 |
| STX16-NPEPL1 | 0.395075302 | 0.04887092 |

| **Table S4** | | |
| --- | --- | --- |
| **Table S4A.** | | |
| Ant-Ins Lowest 20 vs. Highest 20 Psychiatric Morbidity  (Mood Disorders & Controls) | | |
| GeneName | Log2 Fold Change | q-adjusted |
| NPAS4 | -0.12 | 0.02 |
| **Table S4B**. | | |
| Ant-Ins Lowest 20 vs. Highest 20 Psychiatric Morbidity  (Mood Disorders Only) | | |
| GeneName | Log2 Fold Change | q-adjusted |
| No Gene | NS | NS |

| **Table S5** | | |
| --- | --- | --- |
| **Table S5A** | | |
| sgACC Lowest 20 vs. Highest 20 Psychiatric Morbidity (Mood Disorders & Controls) | | |
| GeneName | log2FoldChange | q-adjusted |
| RP1-167G20.1 | -0.877002039 | 0.00452634 |
| RP11-98D18.15 | -0.724926784 | 0.02728451 |
| SNORD113-2 | -0.587366678 | 0.04990453 |
| LAMB3 | -0.444975015 | 0.04653281 |
| C1orf132 | -0.364458633 | 0.03463994 |
| STARD8 | 0.29902623 | 0.04864675 |
| TTC32 | 0.330729284 | 0.045337 |
| FPGT-TNNI3K | 0.411101495 | 0.03670549 |
| TMEM187 | 0.430865111 | 0.02728451 |
| SLC2A4 | 0.455605566 | 0.03670549 |
| VWF | 0.599546082 | 0.04990453 |
| KANK3 | 0.604297074 | 0.01055268 |
| FAM101B | 0.668751171 | 0.04653281 |
| CLDN5 | 0.702514793 | 0.04653281 |
| FLT1 | 0.704786865 | 0.02728451 |
| ADAMTS9-AS1 | 0.741030174 | 0.03670549 |
| ATHL1 | 0.91922823 | 0.04908813 |
| TGFB3 | 0.965307862 | 0.04990453 |
| LRRC32 | 0.96851777 | 0.03124542 |
| TM4SF1 | 1.003397954 | 0.03670549 |
| DDIT4 | 1.040727052 | 0.04990453 |
| CDK2 | 1.099152007 | 0.03670549 |
| IFITM3 | 1.10533146 | 0.04908813 |
| ADAMTS9 | 1.150835433 | 0.03334998 |
| IFITM1 | 1.191864607 | 0.00543532 |
| OSMR | 1.254295112 | 0.04653281 |
| MT1M | 1.284933497 | 0.01055268 |
| IL1R1 | 1.285525005 | 0.04990453 |
| HAMP | 1.290936248 | 0.04908813 |
| KIAA0040 | 1.310680312 | 0.04653281 |
| C10orf10 | 1.548180748 | 0.00378147 |
| ANGPTL4 | 1.560992237 | 0.03463994 |
| STON1-GTF2A1L | 1.599711214 | 0.04908813 |
| MT1X | 1.604506037 | 0.01187363 |
| IFITM2 | 1.681454271 | 0.00441438 |
| SFN | 2.086497044 | 0.03664745 |
| HILPDA | 2.162150261 | 0.00937067 |
| ADM | 2.411503273 | 0.00937067 |
| **Table S5B** | | |
| sgACC Lowest 20 vs Highest 20 Psychiatric Morbidity (Mood Disorders Only) | | |
| GeneName | log2FoldChange | q-adjusted |
| MTND2P28 | -1.474105007 | 0.01310846 |
| ARC | -0.978329496 | 0.02169186 |
| RP13-870H17.3 | -0.844750279 | 0.01707175 |
| RP11-867G23.10 | -0.837706677 | 0.02281121 |
| RP11-52L5.6 | -0.69764366 | 0.02364908 |
| CACNA1G-AS1 | -0.668752739 | 0.01854167 |
| RP1-167G20.1 | -0.602284494 | 0.03793296 |
| DUSP27 | -0.568993179 | 0.04878035 |
| CDC20P1 | -0.540684716 | 0.02889881 |
| F7 | -0.539476544 | 0.00486875 |
| SNORD113-2 | -0.506426759 | 0.01782209 |
| CTB-58E17.5 | -0.491602292 | 0.04391114 |
| RTL1 | -0.482276895 | 0.041726 |
| AC009495.3 | -0.467746838 | 0.04137867 |
| LINC00996 | -0.460621848 | 0.01593218 |
| RP4-778K6.3 | -0.449182168 | 0.03252294 |
| RP4-569M23.4 | -0.416667541 | 0.01746233 |
| LAMB3 | -0.393436452 | 0.02256006 |

**SUPPLEMENTARY FIGURES** [HERE]

**Fig S1. Gene Ontology Terms Derived from WGCNA of Ant-Ins.** **A**, shows Axis-III comorbidity correlated negatively with the Ant-Ins tan module, which is enriched for metabolic, energy transport, mitochondrial translation/gene coexpression genes. **B**, Age at death, Axis-I, and suicide lethality collectively correlated negatively with the yellow module capturing cellular and neuronal ion channel/calcium ion-dependent signaling and synaptic membrane gene coexpression.

**Fig S2. Gene Ontology Terms Derived from WGCNA of Ant-Ins.** **A**, shows Axis-I psychiatric comorbidity being correlated positively with the brown module enriched for a wide-ranging inflammatory cytokine response, T-cell immune response and leukocyte functions gene coexpression in the Ant-Ins, whereas **B**, illustrates the positive correlations between Axis-III comorbidity with the magenta/green module enriched for biosynthesis/protein synthesis and viral transcription in Ant-Ins.

**Fig S3. Gene Ontology Terms Derived from WGCNA of sgACC. A**, shows that sgACC data and identified a positive correlation between Axis-I and the salmon module enriched for spliceosome, thyroid hormone, and notch signaling gene coexpression. B, shows negative correlations between Axis-III comorbidity as well as BMI with the WGCNA tan module enriched for ribosomal, spliceosomal, mRNA transport and methylation, and protein synthesis gene coexpression in sgACC.

**Fig S4. A**, shows Axis-III comorbidity to correlate negatively with the WGCNA grey module capturing cellular immune and developmental regulatory gene coexpression in the sgACC. **B**, shows that the WGCNA red, pink, cyan, tan, grey and green modules enriched for metabolic, protein synthesis and bodily homeostatic regulatory gene coexpression in sgACC were associated with Axis-III comorbidity and BMI.

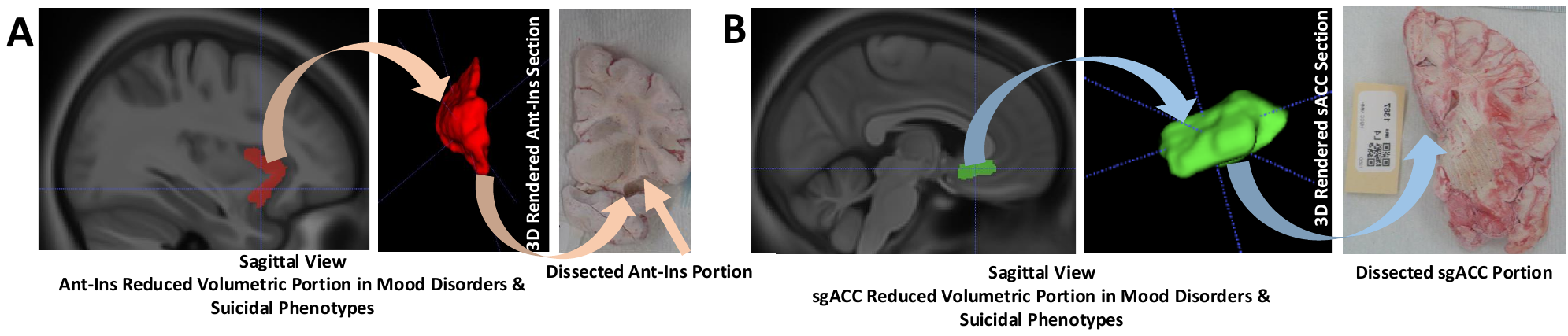

**Fig S5. A**, shows the targeted dissected regional locale of the ventral anterior most portion of the Ant-Ins (**Fig S5A**). Frozen tissue was dissected from the Ant-Ins section for each donor for RNA sequencing by targeting the region most well-documented to be volumetrically reduced in mood disorders. B, shows the dissected regional volume from the sgACC targeting the locale of the ventral ACC Brodmann areas 25 that intersects with the curvature of the Brodmann area 32.

**SUPPLEMENTARY REFERENCES**

<https://www.nimh.nih.gov/health/statistics/suicide.shtml>

<https://www.cdc.gov/injury/wisqars/leadingcauses.html>

<http://www.who.int/mental_health/prevention/suicide/background/en/>

Zaki J. et al. (2016). The Anatomy of Suffering: Understanding the Relationship between Nociceptive and Empathic Pain. *Trends Cogn Sci*, 20(4): 249-259.

Wagner G et al. (2012). Prefrontal cortical thickness in depressed patients with high risk for suicidal behavior. *J Psychiatr Res*. 46(11): 1449-55.

Mathews D. C et al. (2013). Neurobiological aspects of suicide and suicide attempt in bipolar disorder. *Transl Neurosci*. 4(2).

Pan L. A et al. (2013). Differential patterns of activity and functional connectivity in emotion processing neural circuitry to angry and happy faces in adolescents with and without suicide attempt. *Psychol Med*. 43(10): 2129-42.

van Heeringen K., Mann J. J (2014). The neurobiology of suicide. *Lancet Psychiatry*. 1(1): 63-72.

Jollant F et al. (2018). Neuroimaging-informed phenotypes of suicidal behavior: a family history of suicide and the use of violent suicidal means. *Transl Psychiatry*. 8(1): 120.

Eisenberger N. I (2015). Social pain and the brain: controversies, questions, and where to go. *Annu Rev Psychol*, 66: 601-29.

Mee S et al. (2011). Assessment of psychological pain in major depressive episodes. *J Psychiatr Res*, 45(11): 1504-10.

Shneidman ES. (1993). Suicide as psychache. *J Nerv Ment Dis*, 181(3): 145-7.

Strigo IA, Craig AD. (2016). Interoception, homeostatic emotions, and sympathovagal balance. *Philos Trans R Soc Lond B Biol Sci*, 371: (1708).

Collins PY et al. Grand challenges in global mental health. Nature. 2011 Jul 6;475(7354):27-30.

González-Becerra K, Ramos-Lopez O, Barrón-Cabrera E, Riezu-Boj JI, Milagro FI, Martínez-López E, Martínez JA. Fatty acids, epigenetic mechanisms and chronic diseases: a systematic review. Lipids Health Dis. 2019 Oct 15;18(1):178.

Gene B, Dayon L, Kirkland R, Wojcik J, Peyratout G, Severin I, Henry H, Oikonomidi A, Migliavacca E, Bacher M, et al. Blood-brain barrier breakdown, neuroinflammation, and cognitive decline in older adults. *Alzheimer’s & Dementia*, vol. 14, no. 12, December 2018, pp. 1640-1650.

Bettcher B M., Sterling J, Fitch R, Casaletto K, Heffernan K, Asthana S, Zetterberg H, Blennow K, Carlsson C, Neuhaus J, et al. Cerebrospinal Fluid and Plasma Levels of Inflammation Differentially Relate to CNS Markers of Alzheimer’s Disease Pathology and Neuronal Damage. *Journal of Alzheimer’s Disease*, vol. 62, no. 1, February 2018, pp. 385-397.
